# Self-regenerating opsin from reef-building coral

**DOI:** 10.64898/2026.05.24.727572

**Authors:** Yusuke Sakai, Yasushi Imamoto, Shino Inukai, Yuri Tominaga, Tomohiro Sugihara, Takahiro Yamashita, Kota Katayama, Yukiya Kakeyama, Ririka Oka, Erika Okuno, Makoto Iwasaki, Hideki Kandori, Mitsumasa Koyanagi, Akihisa Terakita

## Abstract

Opsins underlie diverse physiological responses to light in animals. In the dark, most opsins bind the chromophore 11-*cis* retinal, which isomerizes to all-*trans* form upon light absorption, representing the initial key step in signaling. Maintenance of opsin function therefore requires continuous regeneration of the inactive, 11-*cis*-retinal-bound state. Here, we report a novel type of opsin, AtAntho2c, from a reef-building coral, whose active form, bound to *all*-trans retinal, can thermally revert to the initial dark state bound to 11-*cis* retinal. A cysteine residue in extracellular loop 2 region plays a key role in the self-regeneration ability. Using time-resolved and low-temperature spectroscopies, we identify two spectrally distinct photointermediates prior to the all-*trans* to 11-*cis* isomerization in AtAntho2c, whose formation rates and yields are found to vary depending on temperature and pH conditions. The active form of AtAntho2c activates Gi/o G protein, resulting in a transient and repeatable decrease in cellular cAMP levels upon repeated light stimulations, even in the absence of exogenous retinal in cultured cells. Furthermore, we confirm that cells expressing AtAntho2c exhibit membrane hyperpolarization via GIRK channel activation light-dependently. These properties highlight the potential of AtAntho2c as a versatile optogenetic actuator capable of repeatedly modulate Gi/o signaling without retinal supplementation.

**Significance Statement:** Light-sensitive proteins, opsins, form active states upon light absorption, leading to intracellular G protein signaling and various cellular outputs. The active states require specific enzymatic machinery or another photon absorption to regenerate inactive opsins ready to respond to repeated light stimuli and maintain continuous responsiveness. In our study, we identify and analyze a coral opsin of which the active state rapidly and autonomously reverts to the inactive state in the dark through thermal isomerization of the retinal chromophore within the opsin. This regeneration mechanism allows the opsin to respond to repeated light stimuli at high temporal resolution and maintain large signal amplitude without the need for exogenous retinal making this opsin potentially useful for developing versatile optogenetic tools.

## Introduction

Animals possess a diversified family of light-sensitive membrane proteins, opsins, which serve as primary photoreceptors for light-dependent physiologies of animals including vision. Animal opsins commonly have the seven transmembrane α-helical structure typical of G protein coupled receptors (GPCRs) and most of them function as light-sensitive GPCRs. In contrast to other GPCRs, opsins uniquely form a covalent bond with a vitamin A-derivative retinaldehyde (retinal) as a light-absorbing chromophore through a conserved lysine residue in the seventh transmembrane helix, forming a retinylidene Schiff base. Most opsins preferentially bind 11-*cis* retinal in the dark to form an inactive dark state. Upon light absorption, 11-*cis* retinal isomerizes to all-*trans* retinal, followed by structural changes of the protein to form an active photoproduct (active state). Namely, opsins specifically bind to 11-*cis* retinal as the inverse agonist or antagonist, and light converts it to the agonist all-*trans* retinal to form the active state. The active state has an ability to interact with heterotrimeric G proteins, activating downstream intracellular signaling and inducing various cell responses (*1*).

Over the past few decades, phylogenetically distinct opsins have been identified across a wide range of species, exhibiting diverse molecular and functional characteristics. Among the various criteria used to classify these opsins, one important feature is the photochemical behavior of their photoproducts (*2*, *3*). The most well-studied opsins, vertebrate visual opsins expressed in vertebrate rod and cone photoreceptor cells, are known as “bleaching opsins” or “monostable opsins” (*2*, *3*) (Fig. 1A). These opsins form active photoproduct metarhodopsin II (meta-II), which has an ability to activate downstream G protein signaling, after isomerization from 11-*cis* to all-*trans* retinal upon light absorption. The photoproduct is unstable and releases the all-*trans* retinal chromophore, which is not followed by the subsequent spontaneous re-isomerization of the retinal to the original 11-*cis* form (*4*) (Fig. 1A). Therefore, to sustain the function of vision, the vertebrate visual system necessitates a metabolic pathway involving a series of enzymes that produces *de novo* 11-*cis* retinal and regenerates dark state pigments, known as the visual cycle (*5*). The other major type of opsins are “bistable opsins” which Gq-coupled invertebrate visual opsins (*6–8*) and some non-visual opsins such as melanopsins (*9*, *10*), parapinopsins (*11*, *12*) and coral Gs-coupled opsins (*13*) are categorized into. Unlike bleaching opsins, bistable opsins produce thermally stable photoproducts, and the chromophore remains bound to the opsin after light absorption. Also, the photoproducts have an ability to revert to the original dark state by absorbing light (Fig. 1B). Because of this property, bistable opsins can regenerate 11-*cis* retinal and restore their original dark state through light-driven reactions, even without requiring specialized enzymatic pathways.

**Fig. 1.**
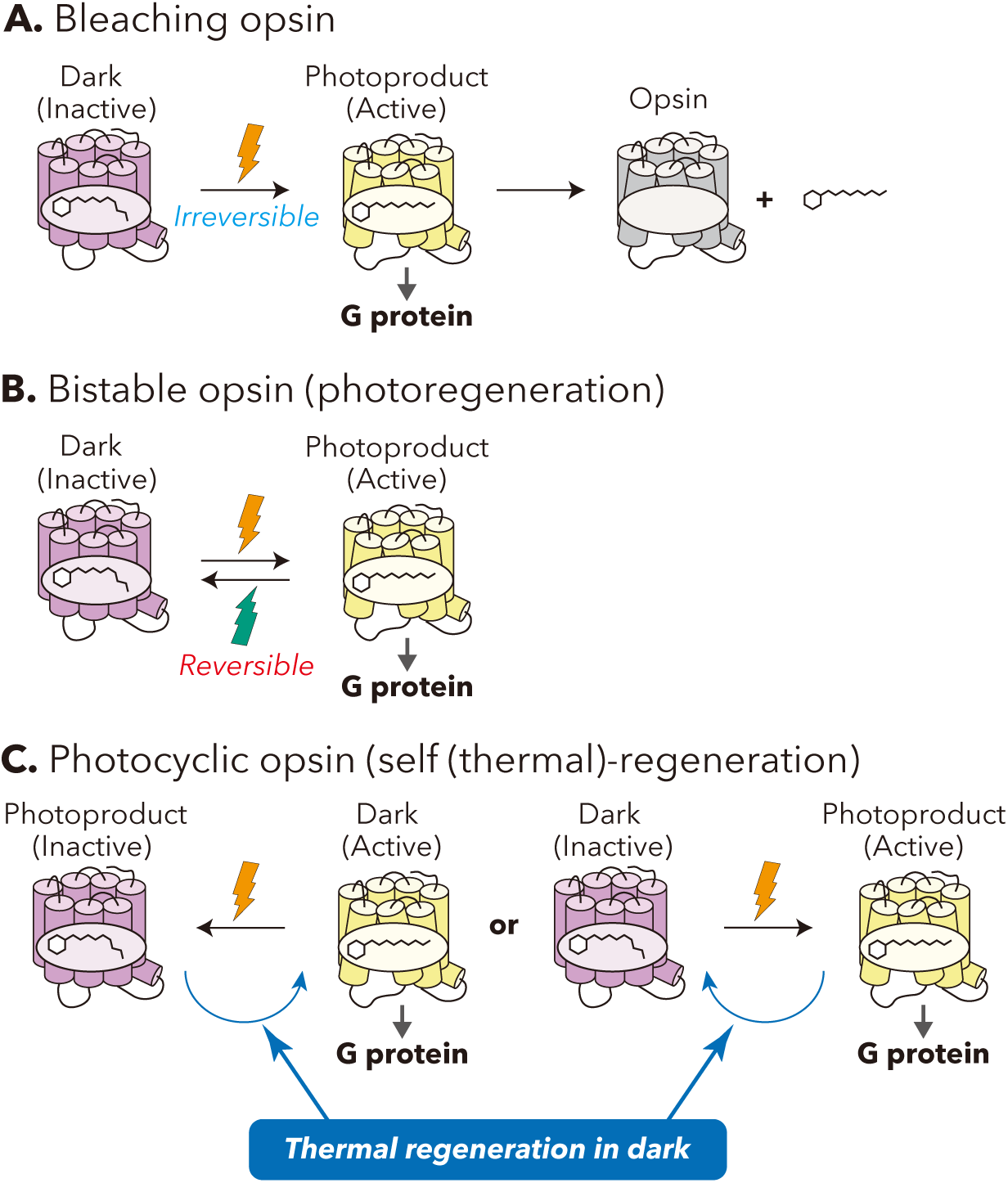
Schematic representation of opsins classified by molecular properties of photoproducts. (A) bleaching opsin, (B) bistable opsin, and (C) photocyclic opsin.

Moreover, recently, a novel type of opsin Opn5L1 has been identified in the chicken genome and represents the first animal opsin having a self-regenerating “photocyclic” nature, a molecular property that enables self-regeneration of the dark state from photoproduct (*14*). Opn5L1 directly forms the active state by binding all-*trans* retinal in the dark, and upon light irradiation, the all-*trans* retinal is isomerized to 11-*cis* retinal to form the inactive state having no or very low G protein activation ability. Notably, Opn5L1 after light absorption (the 11-*cis* retinal-bound form) can thermally revert to the original dark state (the all-*trans*-retinal-bound form) via the formation of a covalent adduct between the retinal chromophore and Cys188 of the opsin protein (Fig. 1C, left). More recently, Sakai et al. (2022) have reported that introduction of a single amino acid mutation, G188C, confers photocyclic nature on the bovine rhodopsin (*15*). The bovine rhodopsin G188C mutant predominantly binds 11-*cis* retinal, and upon light absorption, the chromophore is isomerized to all-*trans* form, leading to meta-II formation. This meta-II then thermally reverts to the original dark state bound to 11-*cis* retinal (*15*) (Fig. 1C, right). In addition, Fujiyabu et al. (2022) have successfully made a photocyclic mutant of mammalian-type Opn5m from *Xenopus tropicalis* by substituting threonine to cysteine at the position 188 (according to bovine rhodopsin amino acid numbering) (*16*). These studies imply that the cysteine residue at position 188 can have a shared significance in the thermal isomerization of chromophore retinal across animal opsins. While many bleaching and bistable opsins have been identified and analysed thus far, only Opn5L1 has been characterized as a natural photocyclic opsin. Consequently, there is little unified understanding of how diverse photocyclic opsins are and to what extent to their underlying mechanisms are shared or diversified.

In this study, we report a novel natural self-regenerating opsin, Antho2c, from a reef-building coral *Acropora tenuis* (hereafter called AtAntho2c), belonging to anthozoan-specific opsin II (ASO-II) group. Unlike the Opn5L1 which binds all-*trans* retinal in the dark and behaves like a retinal receptor, AtAntho2c predominantly binds 11-*cis* retinal in the dark and light irradiation causes the isomerization from 11-*cis* form to all-*trans* form to produce a photoproduct activating G protein signaling cascades as observed in most opsins including vertebrate visual pigments. The all-*trans* retinal-bound active state then thermally reverts to the original dark state bound to 11-*cis* retinal. A series of spectroscopic analyses including a high-speed time-resolved spectroscopy and low-temperature UV-visible spectroscopy allowed us to disentangle the detailed photo-intermediates in the photocycle and also demonstrated that Cys188 of AtAntho2c is important in the photocyclic property. Moreover, our results from the G protein signaling assay showed that AtAntho2c clearly activates Gi/o-type G proteins, triggering subsequent cAMP decrease and change in the membrane potential via GIRK (G protein-activated inwardly rectifying potassium) channel activation. Interestingly, because of the short lifetime of the active state and thermal regeneration of the inactive state, AtAntho2c exhibits transient and repeated cAMP response by consecutive light stimulations. Given these molecular properties, we also discuss the potential of AtAntho2c as an optogenetic tool capable of repeated stimulation with high temporal resolution, particularly its potential for research and therapeutic applications to vision restoration.

## Results

### AtAntho2c shows self-regenerating property demonstrated by spectral and retinal composition changes after light irradiation

AtAntho2c was identified and cloned in the previous study and was expressed in cultured cells to examine its absorption spectrum, where it formed a blue-light-sensitive pigment with a wavelength of maximal absorbance (λ_max_) at 450 nm in the dark (*17*). In this study, to investigate the spectral change and retinal isomerization following light irradiation of wild-type AtAntho2c, we compared the absorption spectra and retinal configurations of the detergent-solubilized pigment before and after irradiation with 460-nm light. However, neither a spectral shift nor retinal isomerization was observed after the light irradiation. The absorption spectrum after the blue-light irradiation completely overlapped with that of the dark state (Fig. S1A). Using high-performance liquid chromatography (HPLC), we showed that AtAntho2c predominantly bound 11-*cis* retinal in the dark and it still bound the same 11-*cis* form even after the 6 min of the blue-light irradiation (Fig. S1B), which is sufficient for most bleaching opsins to complete their photoreaction and for most bistable opsins to achieve photoequilibrium between the dark and light-activated states. On the other hand, we previously found that AtAntho2c induced an increase in intracellular Ca^2+^ level in cultured cells after light stimulation, suggesting that it can light-dependently form a functional active state in the cells (*17*). We therefore hypothesized that, although wild-type AtAntho2c is photoconverted to an active state — spectroscopically distinguishable from the dark state — upon light irradiation, this state thermally reverts to the original dark state in the absence of light (i.e., in the dark) at a rate too rapid to detect either spectral changes or retinal isomerization for our “normal” methods described above. For example, in HPLC analysis, we provided at least 2 min of light irradiation to the purified samples and extracted retinal oximes from the samples more than 4 min after the light irradiation. To test this hypothesis, we employed a high-speed CCD camera spectrophotometer, which allowed millisecond-order recording of time-resolved changes in the absorption spectra before and after irradiation with a < 500-µs yellow (> 490 nm) flashlight. We successfully observed spectral changes of wild-type AtAntho2c following light irradiation. Under conditions of 0°C and pH 6.5, the difference spectra obtained immediately (approximately less than 10 ms) after minus before light irradiation showed a positive peak around 410 nm and a negative peak around 470 nm (Fig. 2A, red curves, black and grey arrowheads), indicating a spectral shift towards shorter wavelengths upon light irradiation. Subsequently, these positive and negative peaks gradually diminished during incubation in the dark, and the difference spectra eventually overlapped with the baseline (Fig. 2A, orange to purple curves). These spectral changes reflect recovery from the blue-shifted spectrum to the original dark spectrum in the absence of light. Next, to elucidate changes in the isomeric composition of the chromophore retinal bound to wild-type AtAntho2c upon light irradiation, we performed HPLC analysis of retinal oxime samples extracted from the purified pigments at five different time points, ranging from immediately (< 5 s) to 10 min after light irradiation. Here, the samples were irradiated for a short period (30 seconds) using a green LED with higher light intensity than that used in Fig. S1B. We observed that 11-*cis* retinal was predominantly bound to the dark state AtAntho2c and was isomerized to all-*trans* retinal, followed by re-isomerization to 11-*cis* form by subsequent incubation for approximately 10 min in the dark at 0°C (Fig. 2B). These data indicate that wild-type AtAntho2c, which binds 11-*cis* retinal in the dark, is converted to the photoproduct bearing all-*trans* retinal and then the photoproduct undergoes thermal self-regeneration to the original dark (11-*cis*-retinal-bound) form upon incubation in the dark, that is, AtAntho2c possesses the self-regenerating photocyclic property.

**Fig. 2.**
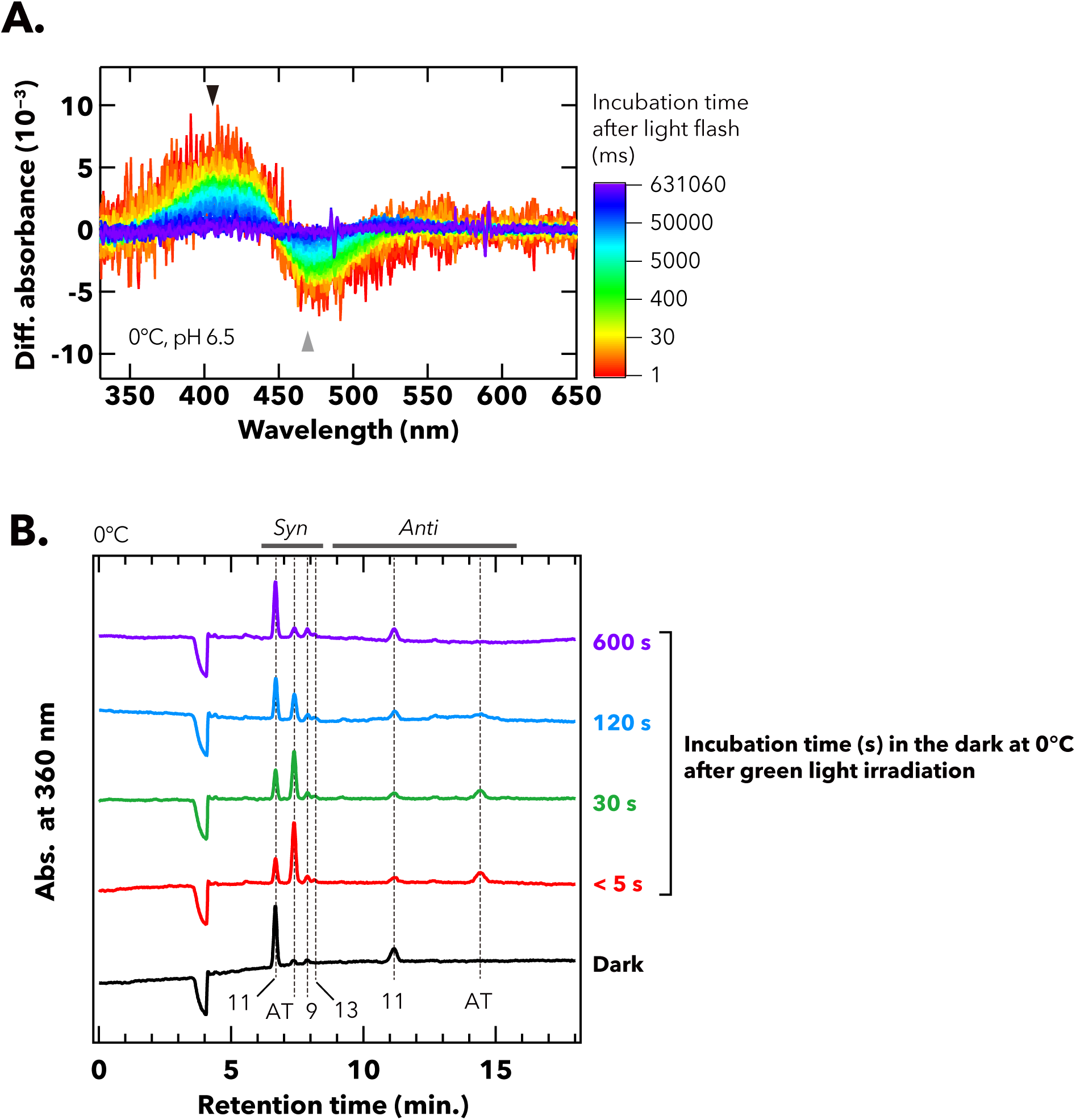
Self-regenerating property of wild-type AtAntho2c. (A) Difference spectra of detergent-solubilized pigment of wild-type AtAntho2c after minus before irradiation with yellow (> 490 nm) flashlight. Spectra were measured using a high-speed CCD camera UV-visible spectrophotometer at 0°C and pH 6.5. Incubation time in the dark after the light irradiation is indicated by a sequential color scheme progressing from red to yellow, green, blue, and purple. Black and grey arrowheads indicate positive peak and negative peaks in the difference spectra at 410 and 470 nm, respectively. (B) The composition of retinal isomers bound to the detergent-solubilized pigment of wild-type AtAntho2c before (“Dark”) and at < 5s, 30 s, 120 s, and 600 s after green-light irradiation. The HPLC was performed after extraction of the retinal from the sample at each time point before and after light irradiation. Samples were kept at 0°C throughout the experiment.

### Cys188 is involved in the self-regenerating property of AtAntho2c

We searched for amino acid residues specific to AtAntho2c that could be involved in the self-regeneration property of the opsin. Notably, AtAntho2c contains a cysteine residue at the amino acid position 188 (according to bovine rhodopsin amino acid numbering) (Fig. 3A). Cys188 has been reported to play a critical role in regulating thermal recovery from the photoproduct to the original dark state in three known photocyclic opsins, Opn5L1 (*14*), the bovine rhodopsin G188C mutant (*15*), and the *Xenopus* Opn5m T188C mutant (*16*). Therefore, we replaced Cys188 with Ser to examine the effect on the self-regenerating property of AtAntho2c. C188S-mutant AtAntho2c formed a blue-sensitive pigment having the λ_max_ at 450 nm in the dark, and the light irradiation induced a substantial shift in the λ_max_ towards shorter wavelengths (Fig. S1C), which was not observed in the wild type (Fig. S1A). Using high-speed spectroscopy, we confirmed that the C188S mutant exhibited a spectral shift to shorter wavelengths just after irradiation with the yellow flashlight as observed in the wild type, but there was no further spectral changes occurred at 0°C and pH 6.5 — the blue-shifted photoproduct of the C188S mutant did not revert to the original dark state after incubation in the dark for 10 min (Fig. 3B). Absorption spectra of C188A-mutant AtAntho2c before and after the yellow-light irradiation were also measured. Although we observed a spectral blue shift after the light irradiation, followed by subsequent spectral changes during dark incubation, no thermal recovery to the original dark state was observed during approximately 150 min of incubation in the dark at 0°C (Fig. S2). HPLC analysis further confirmed that the C188S mutant preferentially bound 11-*cis* retinal in the dark and that this chromophore was isomerized to all-*trans* form immediately (< 5 s) after green-light irradiation. This all-*trans* retinal was not re-isomerized to the 11-*cis* form during the 15 min of dark incubation (Fig. 3C), nor even after 24 h in the dark at 0°C (Fig. 3D). Taken together, these results clearly demonstrate that Cys188 plays an essential role in the self-regeneration process and photocyclic reactions of AtAntho2c, as it does in Opn5L1 (*14*), the bovine rhodopsin G188C mutant (*15*) and the *Xenopus* Opn5m T188C mutant (*16*).

**Fig. 3.**
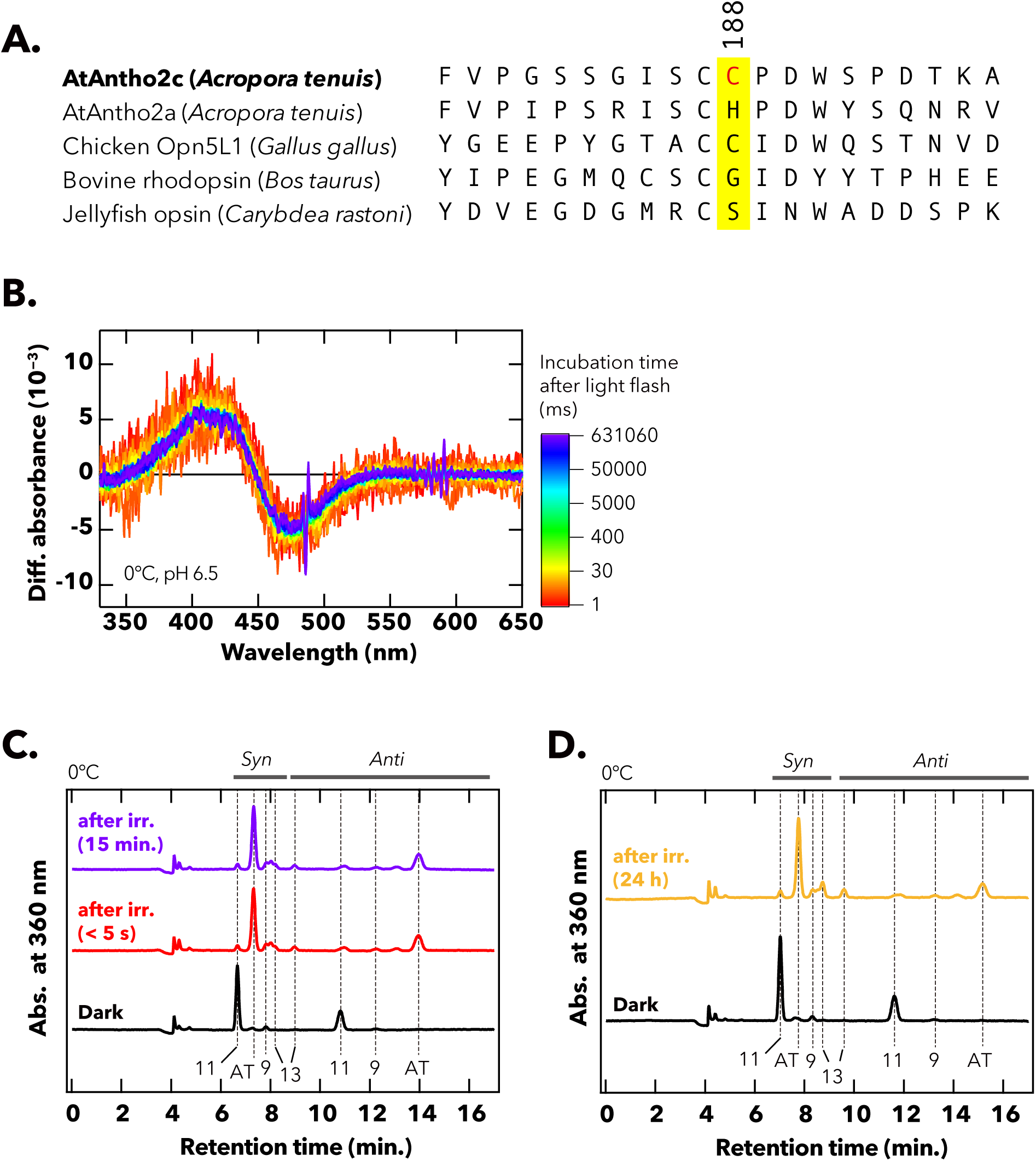
Spectral changes and retinal isomerization of C188S-mutant AtAntho2c upon light irradiation. (A) Partial amino acid sequence of AtAntho2c aligned with other animal opsins. Amino acid residues at the position 188 (bovine rhodopsin numbering) are highlighted in yellow. (B) Difference spectra of detergent-solubilized pigment of C188S-mutant AtAntho2c after minus before irradiation with yellow (> 490 nm) flashlight. Spectra were measured at 0°C and pH 6.5. Incubation time in the dark after the light irradiation is indicated by a sequential color scheme progressing from red to yellow, green, blue, and purple. (C, D) The composition of retinal isomers bound to the detergent-solubilized C188S-mutant AtAntho2c pigment at < 5s and 15 min (C) and 24 h (D) after green-light irradiation, compared with that in the dark (“Dark”, before the irradiation). The HPLC was performed after extraction of the retinal from the sample at each time point before and after light irradiation. Samples were kept at 0°C throughout the experiment.

### The photocycle of AtAntho2c involves spectrally distinct photointermediates generated with temperature- and pH-dependent time constants

We then investigated detailed photointermediates that arise during the photocycle of AtAntho2c. We first measured temporal changes in the absorption spectra of wild-type AtAntho2c after irradiation with yellow (> 490 nm) flashlight, followed by dark incubation under different temperature and pH conditions (Fig. 4). At pH 6.5 and 0°C, wild-type AtAntho2c exhibited a spectral blue shift immediately after irradiation, followed by recovery to the original dark spectrum (Fig. 4A, 0°C, left panel), as also shown in Fig. 2A. At an elevated temperature of 20°C under the same pH condition (pH 6.5), the same initial spectral blue shift was observed immediately after irradiation with the yellow flashlight (Fig. 4A, 20°C, left panel, red curve). Subsequently, a substantial increase in absorbance around 510 nm was observed (highlighted by black arrowhead), indicating a subsequent spectral shift towards longer wavelengths, which was only slightly detected at 0°C and pH 6.5. The spectra eventually overlapped with the baseline (blue to purple curves in Fig. 4A, 20°C, left panel). To capture the time courses of production and decay of the three states — namely, the dark state, the short-wavelength-shifted intermediate, and the long-wavelength-shifted intermediate — we plotted the temporal changes in absorbance at 410, 470, and 525 nm, which are expected to represent transitions among these spectrally different states (middle panels in Fig. 4). The plots showed an increase in absorbance at 525 nm accompanied by a decrease in absorbance at 410 nm within 10^2^ ms at 20°C (Fig. 4A, 20°C, middle panel). They also indicated that the recovery rate to the original dark spectrum (defined as the rate at which the absorbance at 470 nm returns to zero) was faster at 20°C than 0°C under the same pH condition (pH 6.5) (Fig. 4A, middle panel, points versus dashed lines). For further characterization of spectrally distinct photointermediates and the time constants of their formation, UV-visible difference spectra of wild-type AtAntho2c obtained by the high-speed spectroscopy were analyzed using singular value decomposition (SVD) and global fitting methods (Fig. 4, right panels).

**Fig. 4.**
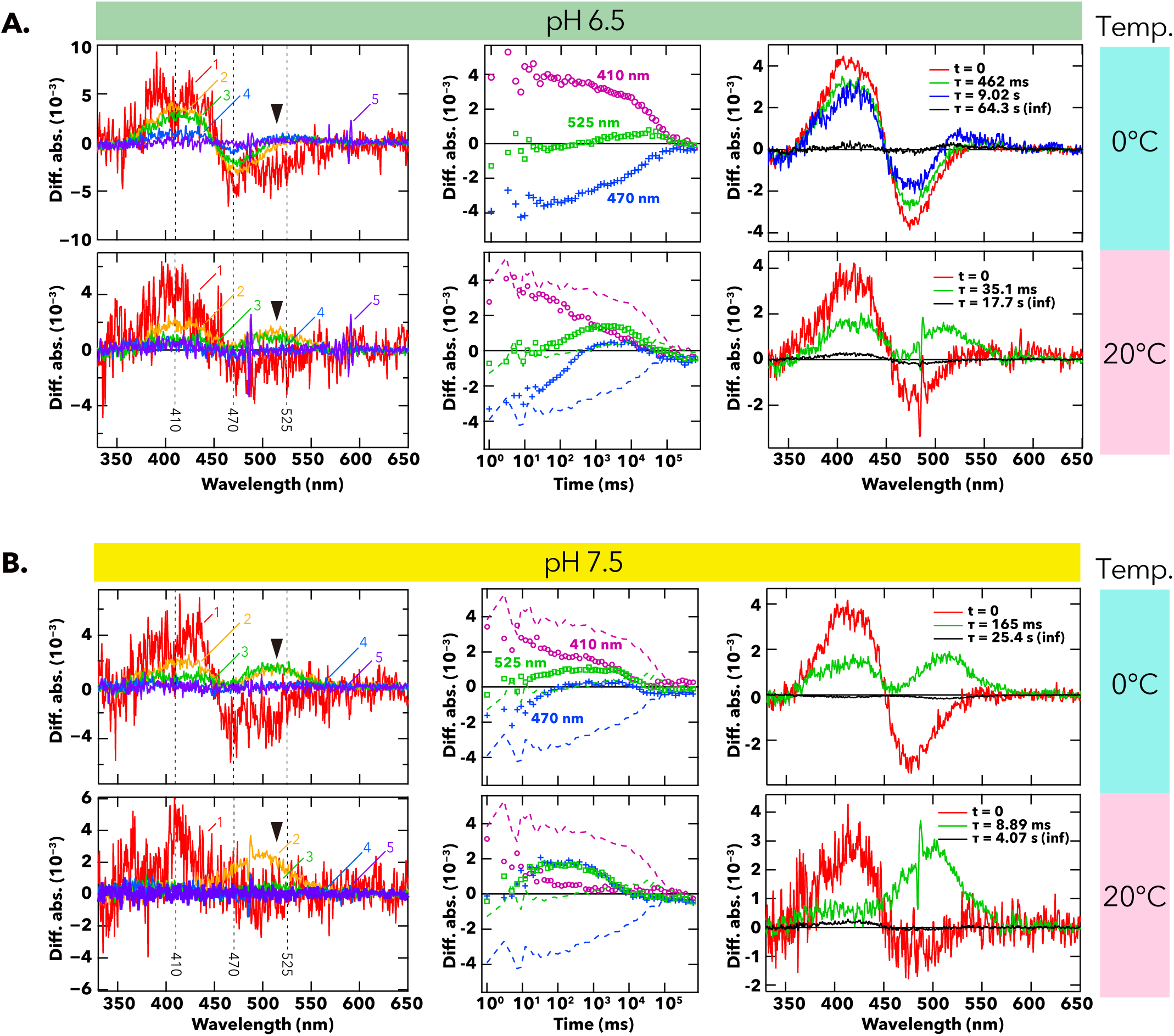
Photoreaction of wild-type AtAntho2c under different pH and temperature conditions. Temporal sequences of spectral changes after yellow-flashlight irradiation to detergent-solubilized pigment of wild-type AtAntho2c were recorded under two different temperatures (0°C and 20°C) at (A) pH 6.5 and (B) pH 7.5. (*left*) Difference spectra of after minus before yellow light irradiation to wild-type AtAntho2c recorded in different time points using a high-speed CCD camera UV-visible spectrophotometer. Colors and curve numbers indicate times of the incubation in the dark after the yellow-light irradiation: 1 (curve 1), 1101 (curve 2), 10101 (curve 3), 100101 (curve 4), and 631060 ms (curve 5). Black arrowheads show positive peaks in the difference spectra around 510 nm. Dotted vertical lines show wavelengths at 410, 470, and 525 nm. (*middle*) Difference absorbance values at 410 (magenta), 470 (blue), and 525 nm (green) are plotted against the time (ms) of incubation in the dark after the light irradiation. Dotted lines in each panel shows the results at pH 6.5 and 0°C reproduced from the top middle panel. (*right*) Spectrally distinct states during photocycle of wild-type AtAntho2c calculated from different spectra shown in left panels using SVD and global fitting methods. The calculated spectra represent specific states after the spectral transition with each time constant.

This analysis allowed the observed spectral changes to be resolved into distinct components corresponding to the states reached after each time-constant-dependent spectral transition. In addition, spectral components extrapolated to time zero (immediately after light irradiation) and to infinite time were also obtained. At 0°C and pH 6.5, wild-type AtAntho2c formed a short-wavelength-shifted intermediate characterized by a positive absorbance peak at ∼410 nm immediately after light irradiation (t = 0; red curve in Fig. 4A, 0°C, right panel). This intermediate was accordingly termed “P_b_”. Following decay of the P_b_ intermediate (τ = 462 ms, green curve), a slight formation of a long-wavelength-shifted intermediate with a positive peak at ∼510 nm (here referred to as “P_r_”) was observed with a time constant (τ) of 9.02 s (blue curve). The spectrum subsequently overlapped with the baseline (τ = 64.3 s; black curve), indicating the final state coincides with the original dark state (Fig. 4A, 0°C, right panel). At the elevated temperature of 20°C at pH 6.5 (Fig. 4A, 20°C, right panel), after generation of P_b_, a substantial amount of P_r_ was generated in the subsequent state with τ = 35.1 ms (green curve), and then the spectrum ultimately returned to the baseline with a time constant of τ = 17.7 s (black curve).

The pH values of the external (seawater) and internal (the gastric cavity) environments of corals, which are in contact with ectodermal and endodermal cells, have been reported to be approximately 8.1 and 6.6 – 8.5, respectively; all these pH values are higher than the pH 6.5 (*18*). To investigate the photochemical reactions of wild-type AtAntho2c under conditions that more closely reflect physiological pH, we recorded light-dependent spectral changes at pH 7.5 (Fig. 4B). Interestingly, spectral changes nearly identical to those observed at the elevated temperature (20°C at pH 6.5) — namely, a large increase in absorbance around 510 nm — were observed at pH 7.5 while maintaining the temperature at 0°C (Fig. 4B, 0°C, left panel). The time-series plot showed an increase in absorbance at 525 nm (and 470 nm), accompanied by a decrease in absorbance at 410 nm (Fig. 4B, 0°C, middle panel), similar to the changes observed under conditions of pH 6.5 and 20°C. Moreover, the rate of recovery to the original dark state under these conditions was faster than that observed at pH 6.5 and 0°C (Fig. 4B, 0°C, middle panel, points versus dashed line). The photochemical processes under pH 7.5 and 0°C calculated by SVD were also similar to those observed at the elevated temperature (20°C) at pH 6.5 (Fig. 4B, 0°C, right panel). A large increase in the P_r_ intermediate was observed after generation of P_b_ with τ of 165 ms (green curve) and both P_r_ and P_b_ subsequently converted back to the original dark state with τ = 25.4 s (black curve). Finally, under conditions of elevated temperature (20°C) and elevated pH (7.5), the difference spectra immediately after yellow-light irradiation showed an increase in the absorbance around 410 nm and a slight decrease around 470 nm (Fig. 4B, 20°C, left panel, red curve). Subsequently, a large increase in absorbance around 510 nm was observed (Fig. 4B, 20°C, orange curve, indicated by black arrowhead), followed by rapid recovery to the original dark state (Fig. 4B, 20°C, left panel, green to purple curves). The time-course plot of the difference absorbance spectra showed that, after a clear increase in absorbance at 525 nm, all measured difference absorbances returned to zero within 10^4^ ms under conditions of elevated temperature and pH (Fig. 4B, 20°C, middle panel), representing the fastest recovery rate among the four pH and temperature conditions tested. SVD analysis (Fig. 4B, 20°C, right panel) showed that, after P_b_ formation, almost all P_b_ was converted to P_r_ with τ = 8.89 ms (green curve), and the spectrum returned to the baseline with τ = 4.07 s (black curve).

Spectral changes of the C188S mutant AtAntho2c were also recorded under different pH and temperature conditions (Fig. 5A, B). While the C188S mutant did not exhibit any subsequent spectral changes during dark incubation after a spectral blue shift by yellow-light irradiation at pH 6.5 and 0°C (Fig. 5A, 0°C), it exhibited distinct spectral changes under conditions of elevated temperature and pH. At the elevated temperature of 20°C, a blue-shifted photoproduct similar to that observed at 0°C was detected immediately after light irradiation, followed by a substantial spectral shift characterized by increases in absorbances at approximately 370 nm and at 510 nm (Fig. 5A, 20°C, left panel; indicated by grey and black arrowheads, respectively). The time-course plot showed that an increase in absorbance at 525 nm occurred concurrently with a decrease in absorbance at 410 nm more than 10^5^ ms after light irradiation (Fig. 5A, 20°C, middle panel). SVD analysis revealed that C188S-mutant AtAntho2c exhibited no spectral transition following the formation of P_b_ at 0°C and pH 6.5 (Fig. 5A, 0°C, right panel). At the elevated temperature of 20°C (pH 6.5), P_b_ was generated immediately after light irradiation and subsequently transitioned to P_r_ and another intermediate with a positive absorbance peak at approximately 370 nm (hereafter referred to as P_uv_) with a time constant of 403.2 s (black curve in Fig. 5A, 20°C, right panel); however, no further reversion to the original dark state was observed. Similar spectral changes to those observed as at 20°C and pH 6.5 were detected at elevated pH (7.5) while maintaining the temperature at 0°C (Fig. 5B). Following light irradiation and subsequent dark incubation, both the P_r_ (∼510-nm peak, black arrowhead) and P_uv_ (∼370-nm peak, grey arrowhead) components were clearly observed in the difference spectra (Fig. 5B, 0°C, left panel). The final spectrum calculated by SVD analysis consisted of two positive peaks and closely resembled that obtained at pH 6.5 and 20 °C (Fig. 5B, 0°C, right panel, black curve). However, the UV-side positive peak was broader, suggesting that it may correspond to a mixed state of P_b_ and P_uv_. Under conditions of both elevated pH (7.5) and elevated temperature (20°C), a more extended sequence of spectral changes, including later-stage transitions, was captured owing to the accelerated spectral dynamics. Under these conditions, yellow-light irradiation initially produced P_b_ (red curve), which subsequently converted to both P_uv_ and P_r_, exhibiting peaks at approximately 370 nm and 510 nm in the difference spectra (blue curve), followed by a gradual conversion of P_r_ to P_uv_ (blue to purple curve, Fig. 5B, 20°C, left panel). Although the C188S mutant exhibited dynamic spectral changes after light irradiation, particularly under conditions of high temperature and pH conditions, these photoproducts did not spectrally convert back to the original dark state. SVD analysis showed that at pH 7.5 and 20°C, the C188S mutant formed P_b_ immediately after light irradiation, which rapidly converted to P_r_ (τ = 12.3 ms, green curve), followed by the accumulation of P_uv_ (τ = 70.3 s, blue curve, Fig. 5B, 20°C, right panel). The final state generated with τ = 392 s consisted predominantly of P_uv_ under these conditions (black curve, Fig. 5B, right panel).

**Fig. 5.**
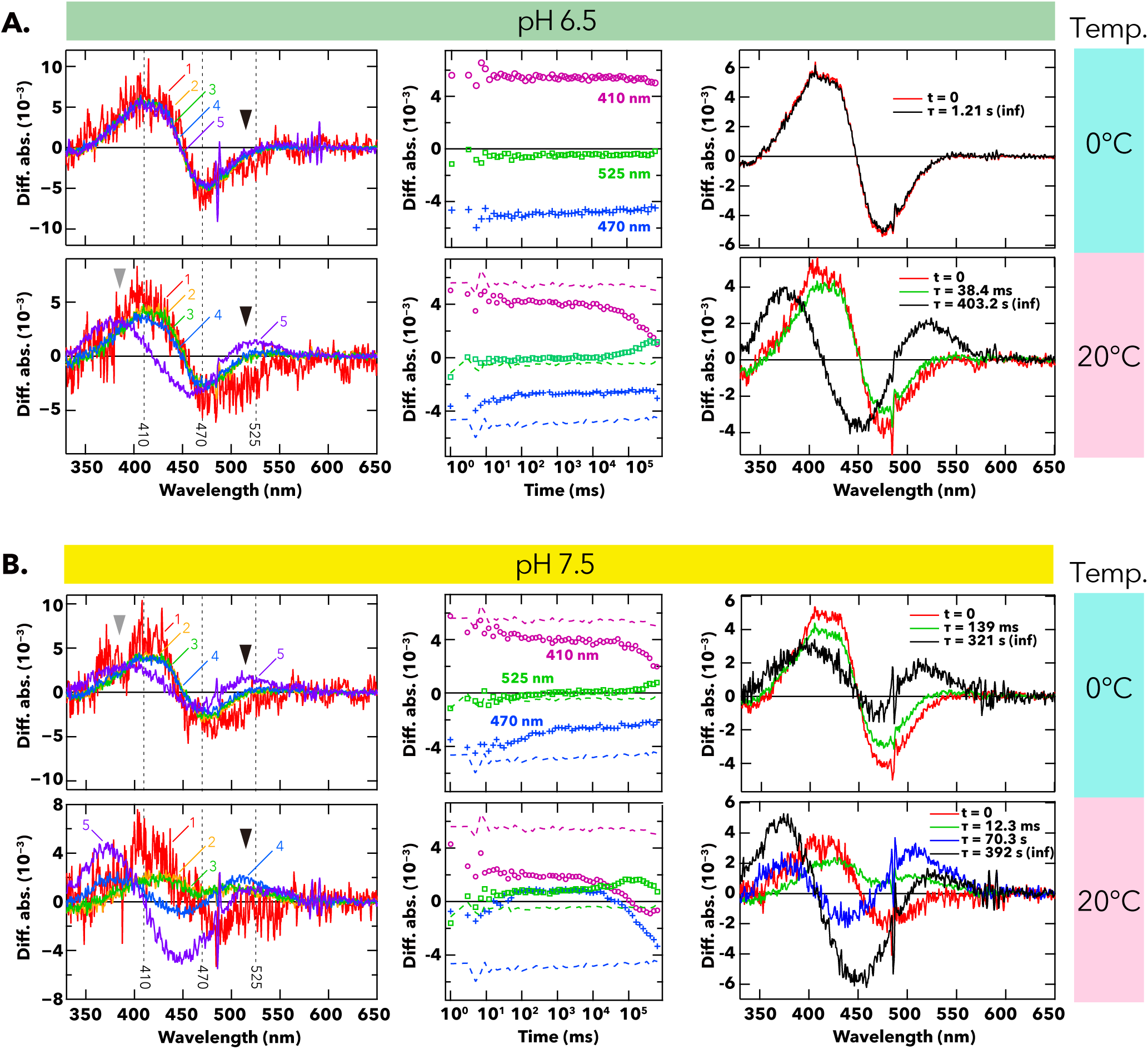
Photoreaction of C188S-mutant AtAntho2c under different pH and temperature conditions. Temporal sequences of spectral changes after yellow-flashlight irradiation to detergent-solubilized pigment of C188S-mutant AtAntho2c were recorded under two different temperatures (0°C and 20°C) at (A) pH 6.5 and (B) pH 7.5. (*left*) Difference spectra of after minus before yellow-light irradiation to C188S-mutant AtAntho2c recorded in different time points using a high-speed CCD camera UV-visible spectrophotometer. Colors and curve numbers indicate times of the incubation in the dark after the yellow light irradiation: 1 (curve 1), 1101 (curve 2), 10101 (curve 3), 100101 (curve 4), and 631060 ms (curve 5). Grey and black arrowheads show positive peaks in the difference spectra around 370 nm and 510 nm, respectively. Dotted vertical lines show wavelengths at 410, 470, and 525 nm. (*middle*) Difference absorbance values at 410 (magenta), 470 (blue), and 525 nm (green) are plotted against the time (ms) of incubation in the dark after the light irradiation. Dotted lines in each panel shows the results at pH 6.5 and 0°C reproduced from the top middle panel. (*right*) Spectrally distinct states during photocycle of C188S-mutant AtAntho2c calculated from different spectra shown in left panels using SVD and global fitting methods. The calculated spectra represent specific states after the spectral transition with each time constant.

### Low-temperature UV-visible spectroscopy captures photointermediates in the AtAntho2c photocycle

To trace the spectral transition observed by high-speed spectroscopy — namely from P_b_ to P_r_ and back to the dark state — starting from an earlier intermediate, we conducted low-temperature UV-visible spectroscopy of wild-type and C188S-mutant AtAntho2c. Specifically, their lipid-reconstituted pigments were irradiated at 77 K, then gradually warmed from 77 K to 290 K in 10 K increments while monitoring the UV-visible spectra at each step. Irradiation of the wild-type sample with 450-nm light at 77 K resulted in the difference spectrum (after minus before light irradiation) showing a large negative peak at ∼460 nm and two positive peaks in the UV (∼397 nm) and green (∼515 nm) regions (Fig. 6A). This early, red-shifted spectrum is considered to indicate generation of an initial Batho intermediate immediately after light irradiation, whereas the increase in UV absorbance may arise from a vibronic band (*19*) or, alternatively, from a partially deprotonated component; however, the origin of this UV absorption cannot be clearly assigned at present. Upon increasing the temperature of the wild-type sample, an intermediate exhibiting a difference absorbance peak around 420 nm was first observed, and its accumulation continued from 90 K to 220 K (Fig. 6A). The blue-shifted intermediate remained detectable over a broad temperature range, from 90 K to 290 K in the difference spectra calculated relative to the dark spectrum as the baseline (positive peaks around 400 nm in Fig. S3A) and was possibly assigned to P_b_ as defined above. As the temperature reached 250 K, we observed an increase and a decrease in the difference absorbances with ∼475-nm and ∼410-nm peaks, respectively, and this spectral change proceeded substantially during incubation at 270 K, corresponding to the conversion of P_b_ back to the original dark state (Fig. 6A). This spectral shift can also be confirmed from the difference spectra calculated relative to the dark spectrum, in which the positive peak around 400 nm (395 – 404 nm) and the negative peak around 470 nm diminished in temperature range from 250 K – 290 K (Fig. S3A). In addition, a substantial increase in the absorbance around 520 nm at 260 K – 270 K was observed in difference spectra relative to the dark spectrum (Fig. S3A), providing evidence of P_r_ generation at high temperature, consistent with the results from high-speed UV-visible spectroscopy shown in Fig. 4. The peak in the range between 500 – 550 nm decreased above 280 K (Fig. 6A). This spectral change indicates that P_r_ reverts to the original dark state upon increasing temperature.

**Fig. 6.**
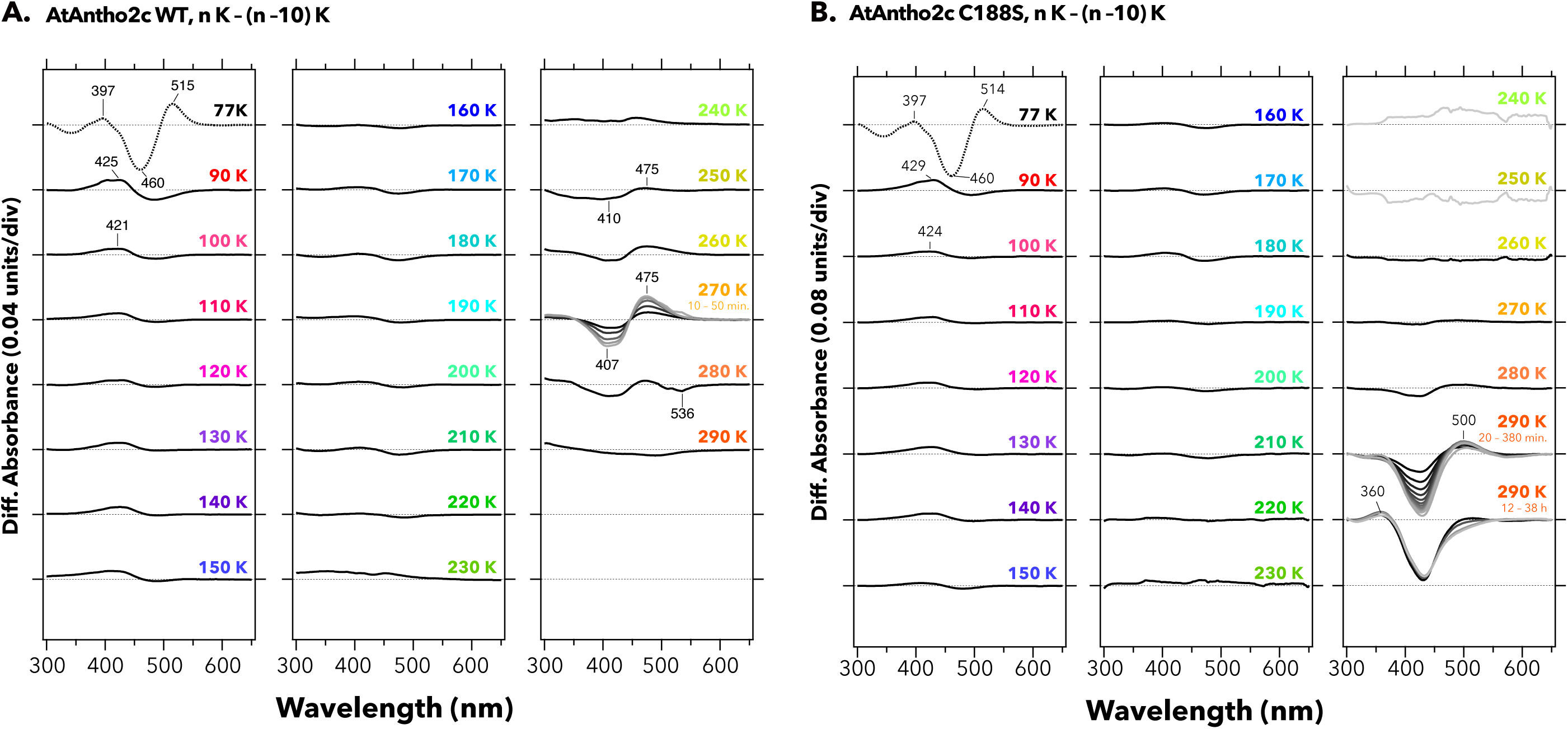
Low-temperature UV-visible spectroscopy and subsequent temperature-dependent spectral changes of (A) wild-type and (B) C188S-mutant AtAntho2c after light irradiation. UV-visible difference spectra after minus before increasing temperature in 10 K increments (n K – (n –10) K) are shown. Lines with different brightness at 270 K (in panel A) and 290 K (in panel B) indicate incubation times at the given temperature (darker to lighter colors correspond to longer incubation).

C188S-mutant AtAntho2c showed a similar initial spectral change to the wild type after 450-nm light irradiation at 77K — a large decrease in absorbance in the blue region and increases in absorbances in the UV and green regions (Fig. 6B, 77K). Following this, the mutant exhibited the generation of P_b_ characterized by a difference absorbance peak around 420 nm, which continued to increase up to ∼210 K (Fig. 6B). P_b_ was detectable from 90 K to 290 K in the difference spectra calculated relative to the dark spectrum (Fig. S3B), similar to the wild type. The formation of P_r,_ characterized by a positive peak at ∼500-nm, began to be observed at 280 K (Fig. 6B). Incubation of the sample at 290 K resulted in a large decrease in the absorbance around 420 nm, indicating the decay process of the P_b_. Simultaneously, the formation of a product showing a difference absorbance peak in UV region was observed, indicating the generation of P_uv_ (Fig. 6B; Fig. S3B), consistent with the results from the high-speed UV-visible spectroscopy (Fig. 5).

In summary, a series of high-speed and low-temperature UV-visible spectroscopic analyses suggests a plausible cycle reaction of AtAntho2c occurring in the dark after light absorption. This photocycle of AtAntho2c comprises the following processes (Fig. 7A): (1) light irradiation of the dark state (D, λ_max_ = ∼450 nm) produces a short-wavelength-shifted intermediate, P_b_, which can be considered the active intermediate, following the production of initial Batho intermediate; (2) the long-wavelength-shifted intermediate, P_r_, characterized by ∼510-nm positive peak in the difference spectra, is generated after P_b_, and is efficiently produced under elevated temperature and pH conditions, suggesting it is in pH equilibrium with P_b_; (3) P_r_ ultimately reverts to the original dark state along with isomerization of the retinal chromophore from all-*trans* to 11-*cis* form, and P_b_ may also directly transition to the original dark state (indicated by dashed line in Fig. 7A). This process may be accelerated under elevated temperature and pH conditions. In contrast to the wild type, the photointermediates P_b_ and P_r_ of the C188S mutant do not revert to the dark state. Instead, the mutant eventually yields mixtures of several intermediates, including P_b_, P_r_ and P_uv_, which probably exist in a state of different equilibrium (Fig. 7B).

**Fig. 7.**
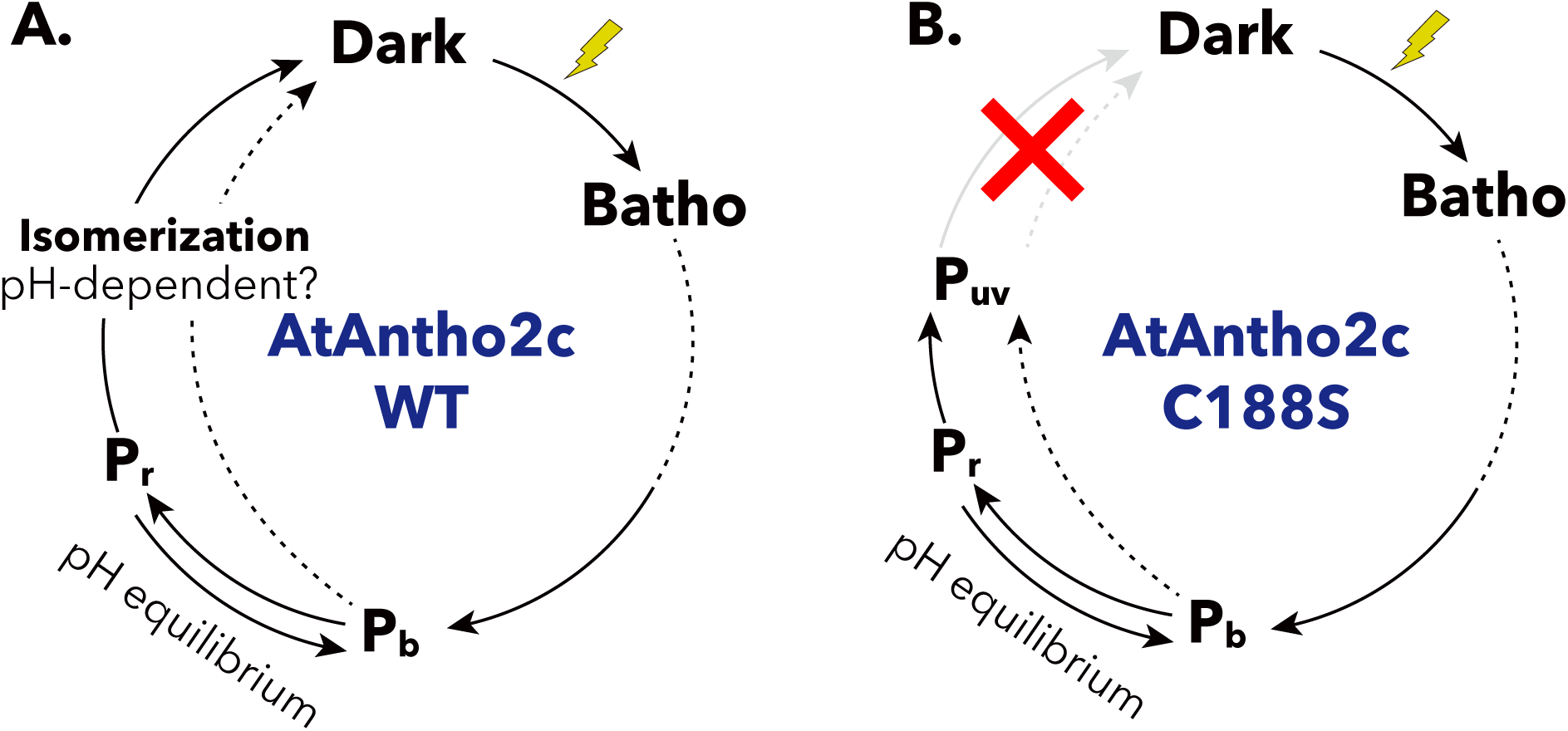
The proposed photocycle scheme of (A) wild-type and (B) C188S-mutant AtAntho2c.

### AtAntho2c activates Gi/o G proteins and induces consistent intracellular cAMP decrease responding to repeated light stimulations

Our previous study reported light-evoked Ca^2+^ elevation in AtAntho2c-expressing cells (*17*). In the present study, we observed a clear transient decrease in cAMP levels in wild-type AtAntho2c-expressing cells after green light illumination (Fig. 8A, black curve), in addition to an increase in Ca^2+^ level (*17*). The cAMP decrease was blocked by the addition of pertussis toxin (PTX), an inhibitor of Gi/o activation (Fig. 8A, red curve), indicating that the decrease in cAMP may be caused by light-induced activation of Gi/o G protein by wild-type AtAntho2c. To directly confirm G protein activation, we performed a NanoBiT G protein dissociation assay, in which dissociation of the Gα and Gβγ subunit upon G protein activation is detected as a decrease in luminescence (*20*) (Fig. 8B). We observed clear transient decreases in luminescence after light irradiation in HEK293S cells co-expressing wild-type AtAntho2c with each of the five individual Gi group G proteins, namely Gαi1, Gαi2, Gαi3, GαoA, and GαoB (Fig. 8C, upper panels). These decreases were statistically significant compared with those observed in no-opsin-expressing cells used as negative controls (Fig. 8C, lower panels). These results show that wild-type AtAntho2c is capable of activating both Gi and Go G proteins.

**Fig. 8.**
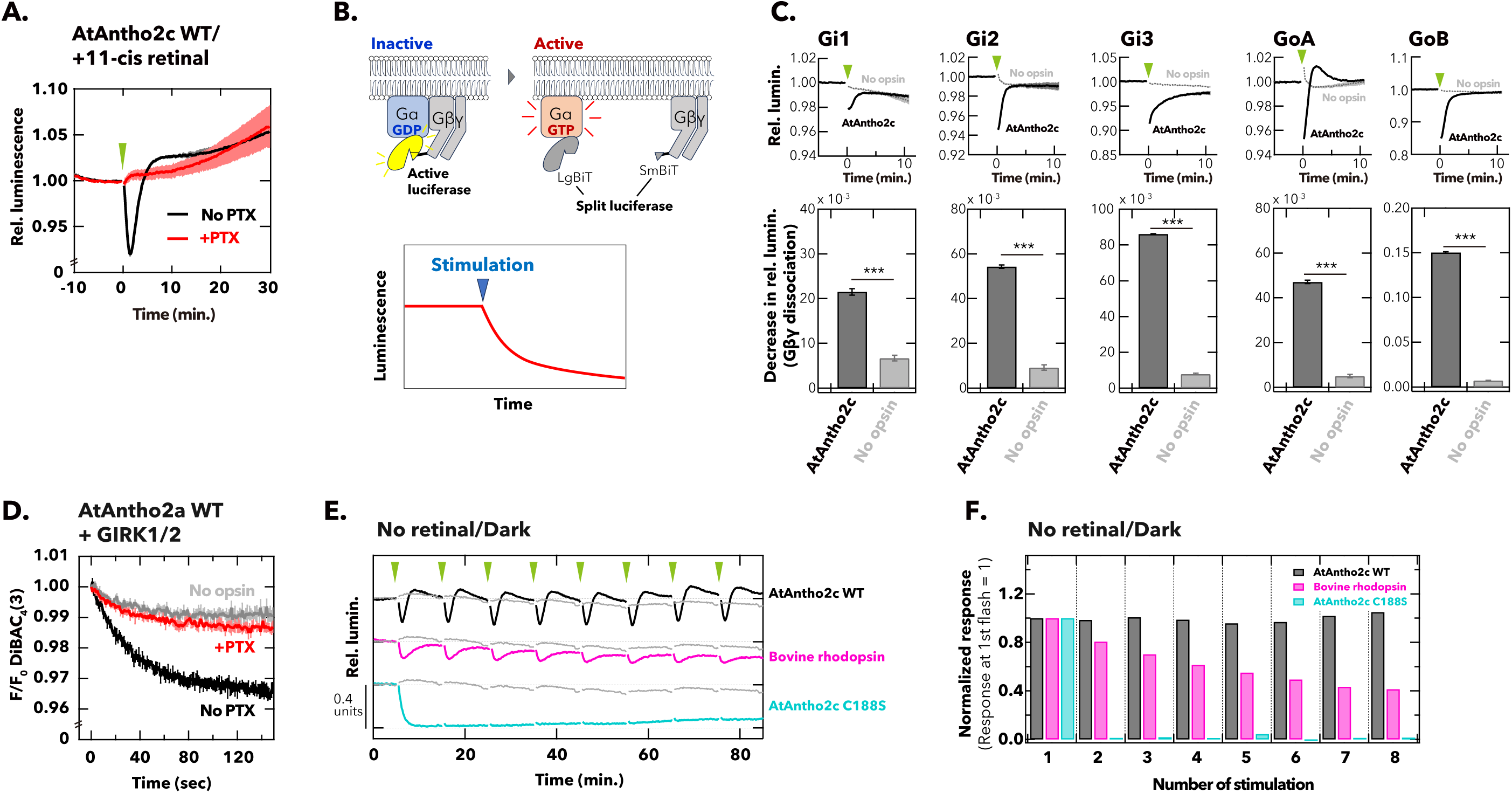
Light-induced G protein activation and downstream signaling in AtAntho2c-expressing cells. (A) Light-induced changes in cAMP level in wild-type AtAntho2c-expressing HEK293S cells. Intracellular cAMP levels were monitored with or without a Gi/o inhibitor pertussis toxin (denoted as PTX in each graph) using GloSensor 22F cAMP sensor. Data are presented as mean (solid lines) ± standard error of the mean (shadings) for n = 3 replicates. Luminescence values were normalized to the average of the 60 s immediately prior to 1-s light irradiation at the time point indicated by a green arrowhead. (B) Schematic of NanoBit G protein dissociation assay. (C) Activation of Gi-group G proteins (Gi1, Gi2, Gi3, GoA, and GoB) by wild-type AtAntho2c assessed using NanoBiT G protein dissociation assay. (*Upper* panels) Time courses of G protein activation levels (Gα-βγ association levels) after light irradiation in AtAntho2c- or no-opsin-expressing HEK293S cells. Data are presented as mean for n = 3 replicates. (*Lower* panels) The maximum decrease in relative luminescence relative to the baseline over 4-min period after light irradiation produced from the raw time-course kinetics of the relative luminescence changes shown in the *upper* panels. Each bar graph with error bar shows the mean ± standard error of the mean for n = 3 replicates. Welch’s t-test was used for pairwise comparison of the results between AtAntho2c-expressing cells and no-opsin-expressing cells (\*\*\**p* < 0.001). (D) Time courses of normalized fluorescence levels from a membrane potential-sensitive fluorescent dye, Bis-(1,3-Dibutylbarbituric Acid) Trimethine Oxonol (DiBAC_4_(3)) in AtAntho2c- (black curves) and no-opsin-expressing cells (grey curve), both co-transfected with GIRK1 and GIRK2. The AtAntho2c expressed in cultured cells was continuously irradiated with 470-nm excitation light for DiBAC4 fluorescence measurement during the assay. The data of the cells treated with pertussis toxin (PTX) is also shown (red curve). Data are presented as mean (solid lines) ± standard error of the mean (shadings) for n = 3 replicates. (E) Time-course kinetics of the cAMP responses to repeated light stimulations in HEK293S cells expressing wild-type AtAntho2c (black), bovine rhodopsin (magenta), and C188S-mutant AtAntho2c. Data from no-opsin-expressing cells were also included as a negative control (grey). Transfected cells were prepared without addition of exogenous retinal and were kept in the dark throughout the entire process prior to the start of recordings. It should be noted that the culture medium contains a small amount of serum-derived retinal. Green arrowheads indicate the time points at which 5-s green-light stimulation was applied. The luminescence values were normalized to the values just before the first light stimulation (time point = 0). (F) Attenuation of response amplitudes across the repeated light stimulation calculated from the data in the panel E. The response amplitude for *N*-th light stimulation was calculated by subtracting the minimum relative luminescence value within the range between *N*-th and (*N+1)*-th light stimulations from the value just before the *N*-th stimulation (baseline). The response amplitude was normalized to the value at the first light stimulation (= 1).

Regarding signaling cascades downstream of Gi/o activation, it is well established that, in addition to Gαi/o-dependent inhibition of adenylyl cyclase and the consequent decrease in intracellular cAMP level, Gβγ dissociated from Gαi/o independently modulates cell excitability by directly regulating the activities of various ion channels, such as G protein-activated inwardly rectifying potassium (GIRK) cannels (*21–23*), voltage-gated calcium (VGCC) channels (*24–26*), and transient receptor potential (TRP) channels (*27*, *28*). In particular, activation of GIRK channels is a key downstream event of the Gi/o G protein signaling pathway, leading to cellular hyperpolarization. Therefore, we next examined whether wild-type AtAntho2c alters cellular excitability via GIRK channel activation after light illumination using a membrane potential-sensitive fluorescent dye, Bis-(1,3-Dibutylbarbituric Acid) trimethine Oxonol (DiBAC_4_(3)). DiBAC_4_(3) enters depolarized cells and increases fluorescence by binding to intracellular proteins and cell membranes. HEK293S cells transfected with wild-type AtAntho2c and GIRK1/2 exhibited a decrease in the DiBAC_4_(3)-derived fluorescence during the measurement compared with no-opsin-expressing cells (Fig. 8D, black versus grey curves), indicating AtAntho2c-induced cellular hyperpolarization. Treatment of the cells with pertussis toxin (PTX) abolished the decrease in the fluorescence (Fig. 8D, red curve). We also confirmed that this decrease in fluorescence was not observed in the absence of the co-expression of GIRK1/2 (Fig. S4). Together, these results indicate that light activation of AtAntho2c induces cellular hyperpolarization via GIRK activation mediated by Gβγ released from activated Gi/o.

The transient decrease in cAMP level after light stimulation and the rapid recovery to the baseline observed in cells expressing wild-type AtAntho2c (Fig. 8A) may reflect its self-regenerating property, in which rapid recovery to the inactive dark state shortened the lifetime of the active state. Given the rapid recovery to the inactive dark state following light stimulation, wild-type AtAntho2c is expected to exhibit cAMP decreases with consistent amplitudes and high temporal resolution, even under rapidly repeated light stimulation. To examine the response stability, we provided repeated light stimulation to AtAntho2c-expressing cells and monitored cAMP responses. As expected, repeated 5-s light stimulation at 10-min intervals elicited repeated decreases in cAMP in wild-type AtAntho2c-expressing cells in the absence of exogenously added retinal (Fig. 8E, black line). We also examined the cAMP response of bovine rhodopsin, which is well-studied bleaching opsin (Fig. 1A) capable of Gi/o G protein activation. Bovine rhodopsin-expression cells showed clear cAMP decreases upon light stimulation in the absence of exogenous retinal, but the response amplitude kept attenuated over the repeated stimulation, possibly because of the pigment bleaching after light absorption (Fig. 8E, magenta line). C188S-mutant AtAntho2c showed a large decrease in cAMP level after the first light stimulation, but in contrast to the wild type, no further decrease in cAMP was observed in response to second and subsequent stimulations under the same condition (Fig. 8E, cyan line). In wild-type AtAntho2c, the response amplitude (the extent of cAMP decrease) was not significantly attenuated across the repeated light stimulation retaining over 90% of the initial response at the 10th light stimulation while bovine rhodopsin showed approximately 35% of the initial response at the last light stimulation and the C188S mutant rapidly diminished the response after the second light stimulation (Fig. 8F). This observation can be attributed to the rapid reversion of the active form to the original dark state in wild-type AtAntho2c, resulting in production of the “ready-to-respond” inactive state even under the continuous or repeated light exposure. Moreover, wild-type AtAntho2c-expressing cells exposed to the room light for 5 h before the recording exhibited maintained large response across the repeated stimulation like the case with the cells without exposure to the room light (Fig. S5).

## Discussion

In this study, we demonstrated a self-regeneration property of AtAntho2c, a coral opsin belonging to the ASO-II group. The inactive dark state bound to 11-*cis* retinal is converted to an active form bound to all-*trans* retinal upon light absorption, which has a potential to activate G proteins, and the active form eventually reverts to the original dark state in the dark via thermal re-isomerization of retinal from all-*trans* to 11-*cis* form. This all-*trans*-to-11-*cis* isomerization is opposite to the 11-*cis*-to-all-*trans* reaction exhibited by Opn5L1, the only native self-regenerating opsin reported to date (*14*), and is similar to the reaction of the bovine rhodopsin G188C mutant, which undergoes thermal isomerization from all-*trans* to 11-*cis* retinal (*15*). In Opn5L1, light-dependent adduct formation via a covalent bond between Cys188 and 11-*cis* retinal disrupts the conjugated double-bond system of retinal (possibly at position C11) and facilitates thermal re-isomerization from 11-*cis* to the all-*trans* form in the dark. During this process, a clear absorbance increase at 270 nm and concomitant decrease in visible region are observed, which likely reflect the disruption of retinal double bounds. In the present study, no clear increase in the absorbance at 270 nm was observed during the dark reaction following light absorption by wild-type AtAntho2c (Fig. S6), similar to the bovine rhodopsin G188C mutant (*15*) and the *Xenopus* Opn5m T188C mutant (*16*). Although no evidence was obtained from the spectroscopic analyses performed here, it remains possible that transient adduct formation between Cys188 and all-*trans* retinal occurs during the thermal recovery of AtAntho2c. In any case, Cys188 appears to be involved in the self-regenerating property shared by AtAntho2c, Opn5L1, bovine rhodopsin G188C and *Xenopus* Opn5m T188C. Despite possible mechanistic differences, these findings suggest that Cys188 functionally involved in the photocycle of phylogenetically divergent animal opsins, while the underlying molecular mechanisms remain to be clarified.

Furthermore, a cysteine residue located close to the retinal chromophore is also significant in the photocycle of a class of microbial rhodopsins, channelrhodopsins. The photocycle of channelrhodopsin-2 (ChR2) is initiated by the absorption of a photon, which drives the isomerization of retinal chromophore from the all-*trans* to the 13-*cis* form. This primary event triggers a cascade of conformational changes that generate a conducting state before the protein eventually returns to its dark state containing all-*trans* retinal (*29*). The kinetics of the photocycle, including the rate at which retinal thermally re-isomerizes from 13-*cis* back to all-*trans*, are strongly influenced by a highly conserved cysteine residue, Cys128, located in transmembrane helix 3 and positioned in close proximity to the C13 atom of retinal. Cys128 forms part of a hydrogen-bonding network, the so-called “DC gate”, together with Asp156 in helix 4 and an intervening water molecule (*30*, *31*). Mutations in the DC gate markedly extend the lifetime of the conducting state by 10^2^–10^5^-fold and slow the photocycle (*32–34*) — these mutational effects have been attributed to alterations in the electrostatic and hydrogen-bonding environments surrounding the retinylidene Schiff base (*31*). In AtAntho2c, as in ChR2, the rate of the photocycle may not be determined solely by Cys188, but rather by interactions such as hydrogen bonds involving Cys188, neighboring amino acid residues, and water molecules. For example, two aspartic acid residues, Asp190 and Asp292, located approximately 7Å from Cys188 in AtAntho2c according to AlphaFold 3 structural model (*35*), are likely candidates for interacting with Cys188 and modulating the photocycle rate. The pH-dependent changes in the photocycle rate observed in AtAntho2c further suggest that protonation states of titratable amino acids, and possibly the Schiff base in the P_b_ and P_r_ intermediates, are crucial for the reactions. In ChR2, a sequence of deprotonation and re-protonation events of the Schiff base, in which the DC gate plays a central role, is rate-limiting for the photocycle, and consequently the photocycle kinetics are pH-dependent (*36*). Our low-temperature spectroscopy revealed spectral features in the Batho intermediate that may suggest partial deprotonation of the Schiff base in AtAntho2c (Fig. 6A), although this assignment cannot be made conclusively at present. If such a state is indeed formed, it could reflect the rapid proton transfer involving the Schiff base and nearby potentially titratable residues immediately after light absorption. We have previously revealed that an AtAntho2 opsin, which belongs to the same ASO-II group, employs chloride counterion to stabilize the protonated Schiff base in the dark state and switches it to Glu292 in the photoproduct (*17*). AtAntho2c also had an acidic amino acid Asp292, and thus if the same mechanism applies to AtAntho2c, the spectral features observed in the Batho intermediate might be associated with a transient change in counterion interactions, such as a switch from chloride ion to Asp292; however, further experimental evidence will be required to establish this possibility. A detailed description of how the interactions among Cys188, the Schiff base, and other potential amino acid residues (such as Asp292) vary as a function of pH may, in future studies, provide important insights into the molecular mechanism underlying the AtAntho2c photocycle.

Recently, animal opsins have attracted great attention as template upon which to develop optogenetic tools that can regulate intracellular G protein signaling (*37–42*). In this study, we demonstrate that AtAntho2c drives intracellular Gi/o signaling cascades in a light-dependent manner in a cultured cell system as shown in Fig. 8. Owing to its self-regenerating property, AtAntho2c exhibits rapid and consistent responses to repeated light stimulation. These responses were observed even in the absence of exogenously added retinal (Fig. 8E; Fig. S5), indicating that AtAntho2c achieves functionality by utilizing a small amount of retinal from serum-derived vitamin A metabolites present in the culture medium. These molecular features of AtAntho2c may be advantageous for the development of GPCR optogenetic tools that provide stable response amplitudes with high temporal resolution, even under a limited supply of 11-*cis* retinal. For example, AtAntho2c has potential as an optogenetic actuator for visual restoration. Inherited retinal degenerations, such as retinitis pigmentosa (RP), lead to loss of photoreceptor cells, sever visual impairment, and, in the most advanced cases, complete blindness (*43*, *44*). Optogenetic strategies for visual restoration aim to confer light sensitivity on surviving inner retinal neurons, including bipolar cells and retinal ganglion cells, by ectopically expressing light-sensitive proteins and thereby converting them into photoreceptor-like cells (*44–46*). Among these targets, ON-bipolar cells (OBCs) express the metabotropic glutamate receptor 6 (mGluR6), which activates the Go signaling cascade upon receiving photoreceptor input (*47*) and, in turn, closes TRPM1 non-selective cation channels, leading to cellular hyperpolarization (*48*, *49*). Therefore, therapeutic optogenetic approaches that introduce Go-coupled animal opsins into OBCs are expected to restore visual sensitivity by engaging the native mGluR6 downstream signaling pathway in OBCs. Multiple optogenetic actuators engineered from bleaching or bistable animal opsins (Fig. 1A and B) have been ectopically expressed in OBCs and have been shown to restore visual function at both electrophysiological and behavioral levels in *rd1* mice, a model of photoreceptor degeneration (*39*, *50–53*). One potential limitation of this approach is a significant attenuation of the light response upon repeated stimulation, particularly with bleaching opsin-based tool(*51*). AtAntho2c identified in this study natively activates the Go signalling cascade (Fig. 8C) and, owing to its self-regeneration capacity, is expected to maintain consistent activation in response to repeated light stimulation without significant attenuation of signal intensity (Fig. 8E, F). These properties suggest that AtAntho2c could support robust cellular responses with high temporal precision under continuous light exposure. In addition, AtAntho2c retains its bound retinal chromophore after photoactivation rather than releasing *all-trans* retinal. This property may confer an important advantage for visual restoration therapies, as it minimizes the release of free all-*trans*-retinal of which accumulation has been implicated in photoreceptor and retinal toxicity (*54–56*). Together with its ease of use, stemming from the ability to retain functionality even under the room light condition (Fig. S5), AtAntho2c represents a highly promising optogenetic tool for both research and therapeutic applications to visual restoration.

The physiological role of AtAntho2c in corals remains unknown. One possible function is that it modulates the symbiotic relationship between corals and photosynthetic unicellular algae (dinoflagellates). A single-cell RNA sequencing (scRNA-seq) analysis of the reef-building coral *Stylophora pistillata*, revealed that an opsin is specifically expressed in a distinct population of endodermal cells, namely alga-hosting cells, which exhibit 50% – 83% algal occupancy (*57*). Here, based on the scRNA-seq database publicly provided by the authors (https://sebe-lab.shinyapps.io/Stylophora_cell_atlas/), we identified this opsin as a homologue of AtAntho2c that possesses the conserved Cys188 residue. This finding suggests that Antho2c may be involved in regulating light-dependent physiological processes in host cells, such as controlling cellular transcriptomic and metabolic states in response to algal photosynthesis. Thus, investigating the coordination of host-cell physiology with symbiont photosynthesis, with a focus on Antho2c homologs, may provide new insights into the mechanisms underlying coral-algal photosymbiosis.

## Materials and methods

### Construction of expression vectors

The cDNA of AtAntho2c was cloned as described in the previous study (*17*). Briefly, the open reading frame of wild-type AtAntho2c was tagged with rho 1D4 epitope sequences (ETSQVAPA) at the C-terminus. Site-directed mutants were produced by overlap extension PCR using PrimeSTAR Max DNA Polymerase (TAKARA, Shiga, Japan) using site-specific primers and were also tagged with the rho 1D4 epitope sequence. The tagged cDNAs were inserted into the HindIII/EcoRI-digested pUSRα vector (*58*) or the EcoRI/NotI-digested pMT vector (*59*) using In-Fusion HD cloning kit (TAKARA).

### Opsin expression in cultured cells and UV-visible spectroscopy

The plasmids containing ORFs of wild-type and C188S-mutant of AtAntho2c (15 µg per 100 mm culture dish) were transfected into COS-1 cells by the polyethyleneimine (PEI) transfection method (*60*, *61*). The transfected cells were incubated for 24 h at 37°C under 5% CO_2_. After addition of 11-*cis* retinal (1 µL of 4 mM 11-*cis* retinal per 100 mm culture dish), the cells were incubated another 24 h at 30°C in the dark until collecting the cells. The reconstituted pigments were extracted from the cell membranes in the buffer containing 1% dodecyl β-D-maltoside (DDM, Dojindo, Kumamoto, Japan), 50 mM HEPES, and 140 mM NaCl (pH 6.5). The detergent-solubilized samples were mixed with 1D4-condjugated agarose beads overnight, and the mixture was washed on Bio-Spin columns (Bio-Rad, Hercules, CA, USA) using the buffer containing 0.02% DDM, 50 mM HEPES, and 140 mM NaCl (pH 6.5, buffer A). The purified pigments were eluted with buffer A containing 0.5 ∼ 1 mg/mL 1D4 peptide (custom peptide synthesis by GenScript Japan Inc., Tokyo, Japan). UV-visible spectroscopic analyses were performed using a V-750 UV-visible spectrophotometer (JASCO Corporation, Tokyo, Japan). For each spectral measurement, the absorbance spectrum was recorded from 250 to 750 nm at a scanning rate of 400 nm/min, and three consecutive scans were performed for the same sample. A 100-W halogen lamp was equipped on the spectrophotometer and used to illuminate samples with a set of optical interference filters (450 nm, Toshiba, Tokyo, Japan) or a cutoff filter (Y-49, Toshiba).

### Retinal configuration analyses by high performance liquid chromatography (HPLC)

Purified pigments of wild-type and C188S-mutant AtAntho2c were placed on ice and then were kept in the dark or illuminated for 30 s with green scope LED light (color: “Cyan”, Ex-DHC; BioTools Inc., Gunma, Japan). Retinal chromophore bound to the samples in the dark and at different time points after the green light irradiation were extracted as described previously (62), with some modifications. Briefly, 100 µL of samples were mixed with 210 µL of −20°C 90% methanol and 30 µL of 1 M hydroxylamine to convert retinal chromophore in a sample into retinal oxime. The retinal oxime was extracted with 700 µL of n-hexane. 200 µL of the extract were injected into a YMC-Pack SIL column (particle size 3 μm, 150 × 6.0 mm) and eluted with n-hexane containing 15% (v/v) ethyl acetate and 0.15% (v/v) ethanol at a flow rate of 1 mL/min.

### Time-resolved UV-visible spectroscopy using a high-speed CCD camera spectrophotometer

Absorption spectra were measured using a high-speed CCD camera spectrophotometer (C10000 system, Hamamatsu Photonics, Shizuoka, Japan) in the dark and at different time points after light irradiation. The flashlight was generated from a Xenon flash lamp (SA-200, Nissin Electronic; pulse duration of ∼170 µs and flash lamp input of 200 J/F) and was passed through a cutoff filter (Y-49, AGC Techno Glass Co., Shizuoka, Japan). The temperature of the sample was kept at 0°C or 20°C by a temperature controller (pqod, QUANTUM Northwest, Liberty Lake, WA, USA). The pH of the samples was adjusted with 100 mM CAPS containing NaOH. pH values were measured using a pH meter (B-211; HORIBA, Kyoto, Japan) immediately after each spectroscopic measurement.

### Singular value decomposition (SVD) and global fitting analysis

The transient difference spectra after yellow flash excitation were analyzed by singular value decomposition (SVD) method using Igor Pro 6 (WaveMetrics, Portland, OR, USA) as reported previously (63). The matrix 𝐀 containing the series of transient difference spectra was decomposed as follows:

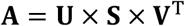

Significant 2 – 4 columns of 𝐕^T^ were globally fitted by the sum of exponential functions to determine time constant (1/*k*) and decay-associated difference spectrum (B spectrum) of each transition. The extrapolated difference spectra at *t* = 0 and infinite time, and that of the pseudo-steady state after each transition are presented.

### Opsin expression in insect cells and purificaction

The cDNAs encoding wild-type and C188S-mutant AtAntho2c were subcloned from the pUSRα vector into the pFastBac HT vector (Bac-to-Bac Baculovirus Expression System; Thermo Fisher Scientific, Waltham, MA, USA) using the In-Fusion cloning kit (TAKARA). For the AtAntho2c construct, a human rhinovirus (HRV) 3C protease cleavage site was introduced at the C-terminus, followed by GFP and 8×His tag to facilitate protein purification. Protein expression was carried out as described previously (64). Briefly, recombinant baculoviruses were used to infect *Spodoptera frugiperda* (Sf9) insect cells, and membrane fractions were prepared from harvested cells. The purified membrane samples were reconstituted overnight with 11-*cis*-retinal at a final concentration of 30 μM. Reconstituted pigments were extracted from the cell membranes in the buffer containing 1% (w/v) DDM, 0.2% (w/v) Cholesteryl hemisuccinate (CHS, Sigma, St. Louis, MO, USA), 30 mM HEPES, and 500 mM NaCl (pH 7.0). The detergent-solubilized samples were mixed with TALON metal affinity resin (TAKARA) for 2 h at 4°C and subsequently washed with buffer containing 0.1% DDM, 0.02% CHS, 30 mM HEPES, 500 mM NaCl, and 3 mM imidazole (pH 7.0). Bound proteins were eluted with buffer containing 0.1% DDM, 0.02% CHS, 30 mM HEPES, 500 mM NaCl, and 350 mM imidazole (pH 7.0). The eluted-samples were further purified with GFP nanobody-conjugated agarose resin (GFPNb resin) for 2 h at 4°C, followed by washing with buffer containing 0.01% DDM, 0.002% CHS, 20 mM HEPES, and 140 mM NaCl (pH 7.0). The target protein was eluted by adding HRV 3C protease (TAKARA). Purified AtAntho2c pigments were reconstituted into phosphatidylcholine (PC) liposomes using buffer containing 50 mM HEPES, 140 mM NaCl, 0.75% CHAPS, and 1 mg/ml PC at a protein–to–lipid molar ratio of 1:30. Dialysis was performed in order to remove detergent and the reconstituted samples were finally suspended in buffer containing 2 mM NaH_2_PO_4_ and 10 mM NaCl (pH 7.0).

### Low-temperature UV-visible spectroscopy

Reconstituted pigments of wild-type and C188S mutant AtAntho2c were placed on a BaF_2_ window (Pier Optics Co., Ltd., Gunma, Japan) and allowed to air-dry overnight at 4°C, followed by additional drying under reduced pressure using an aspirator. For hydration, approximately 1.0 µL of H_2_O was placed next to the dried film, and the sample was sealed with another BaF_2_ window using a silicon rubber O-ring, resulting in an optical path length of 0.5 mm. Low-temperature UV-visible spectroscopic analysis were performed using a V-750 UV-visible spectrophotometer (JASCO). Measurements were performed on hydrated films mounted in a cryostat (OptistatDN; Oxford Instruments, Oxford, UK) equipped with a temperature controller (ITC4; Oxford Instruments, Oxford, UK), with liquid nitrogen used as the coolant. The light source was a 1 kW halogen–tungsten lamp (Master HILUX-HR; Rikagaku, Tokyo, Japan). To generate the initial Batho intermediate, samples were illuminated with 450-nm interference filters (03-FIV-004; Melles Griot, Carlsbad, CA, USA) for 5 min at 77 K. Changes in the λ_max_ values were measured by gradually warming the photoproducts from 77 K to 290 K in approximately 10 K increments. At each temperature step, samples were equilibrated for 20 min prior to spectral acquisition.

### GloSensor cAMP assay

cAMP levels in opsin-expressing cultured cells were assessed by GloSensor cAMP assay (Promega, Madison, WI, USA) as described previously (65–67). Briefly, HEK293S cells seeded in 35-mm dish were transfected with 1.5 µg of the plasmid containing open reading flames of each opsin with 1.5 µg of pGloSensor-22F cAMP plasmid (Promega) using the polyethyleneimine (PEI) transfection method. The transfected cells were incubated for ∼24 h at 37°C under 5% CO_2_ with adding 0.2 µM/dish of 11-*cis* retinal 4 – 5 h after the transfection. The cultured medium was replaced with a medium containing GloSensor cAMP Reagent Stock Solution (Promega), and the cells were incubated to equilibrate with the media at 25°C for at least 2 h. After recording baseline in the dark, the cells were stimulated with green light (color: ‘Cyan’, Ex-DHC, BioTools Inc, Gunma, Japan) and luminescence values were recorded using GloMax 20/20n Luminometer (Promega). For the inhibition of Gi/o G protein activation, pertussis toxin (PTX) (FUJIFILM Wako Pure Chemical Co., Osaka, Japan) diluted in a buffer containing 10 mM potassium phosphate, 137 mM NaCl, and 10% glycerol (final concentration of 200 ng/mL) was added twice (just before the addition of 11-*cis* retinal and at the timing of replacing the culture medium 2-h before starting each measurement). For repeated light stimulation tests, the transfected cells (each opsin and pGloSensor-22F plasmids) were treated in the same way as described above but 11-*cis* retinal was not added. The cultured medium was replaced with a medium containing GloSensor cAMP Reagent Stock Solution (Promega), and the cells were incubated at 25°C for 5 h in the dark (for Fig. 8E, F) or exposed to the room light condition for 5 h (for Fig. S5A, B) prior to the start of the recordings.

### NanoBiT G protein dissociation assay

G protein activations by wild-type and C188S-mutant AtAntho2c were investigated in HEK293S cells using the NanoBiT-G-protein dissociation assay (*20*). HEK293S cells in 35-mm dish were transfected with 2.4 μg of each opsin plasmid, 0.06 μg of Gα-LgBiT, 0.3 μg of Gβ1-SmBiT, and 0.3 μg of Gγ2 plasmids using the PEI transfection method, as described in a previous report (67, 68). The transfected cells were incubated overnight at 37°C. Before measurements, the culture medium was replaced with a CO_2_-independent medium containing 10% FBS, Nano-Glo Vivazine Substrate (Promega), and 11-*cis* retinal, and the cells were incubated for at least 4 h to equilibrate with the media. Luminescence values were measured at 25℃ using a GloMAX 20/20n Luminometer (Promega). The cells were illuminated with green LED light (495 nm) for 5 s. The G-alpha subunit sequences used for the Gα-LgBiT constructs are as follows; rat Gαi1 (NP_037277.1), rat Gαi2 (NP_112297.1), rat Gαi3 (NP_037238.1), mouse GαoB (NP_001106855.1), and monkey GαoA (NP_001182358.1). The G-beta subunit sequence used for the Gβ1-SmBiT construct was derived from human Gβ1 (NP_001269468.1), and the G-gamma subunit sequence was derived from human Gγ2 (NP_001230702.1). The GαoA-LgBiT, Gβ1-SmBiT, and Gγ2 plasmids were kindly provided by Dr. Takashi Nagata and Dr. Keiichi Inoue (The University of Tokyo) (68). The other Gα-LgBiTs were constructed based on the nucleotide sequence of GαoA-LgBiT.

### Measurement of membrane potential change

We monitored membrane potential changes of the cells expressing AtAntho2c using a fluorescent dye, Bis-(1,3-Dibutylbarbituric Acid) Trimethine Oxonol (DiBAC_4_(3), Dojindo, Kumamoto, Japan). DiBAC_4_(3) added to the extracellular medium exhibits a membrane potential–dependent distribution across the plasma membrane. Upon depolarization of the plasma membrane, increased influx of the dye into the cytoplasm occurs, where the internalized dye produces an enhancement of fluorescence intensity. HEK293S cells seeded in 35-mm dish were transfected with 2.4 µg of opsin plasmid, 1.5 µg of GIRK1, 1.5 µg of GIRK2, and 0.225 µg of pRSVTAg using the PEI transfection method. After incubating the transfected cells for 4 – 5 hours at 37°C, 0.2 µM/dish of 11-*cis* retinal was added to the cells and further incubated under 37°C and 5% CO_2_ condition overnight. The culture medium was washed twice with a CO2-indeoendent assay medium (20 mM HEPES, 120 mM NaCl, 2 mM KCl, 2 mM CaCl_2_, 1 mM MgCl_2_, 5 mM Glucose, and 5 µM DiBAC_4_(3), pH 7.4) and then the cells resuspended in 1 mL of the assay medium were seeded in 96-well plate (100 µL/well). After 30-minute-incubation at 37°C and 5% CO_2_, fluorescence values from each well were recorded under 37°C using a plate reader (Optima FLUOStar, BMG Labtech, Ortenberg, Germany) with an excitation wavelength of 485 nm and an emission wavelength of 520 nm (1 s exposure). The excitation light was also used for stimulating the cells expressing opsin. Fluorescence recordings were sampled at 1-s temporal resolution. PTX (FUJIFILM Wako Pure Chemical Co., Osaka, Japan) was added for the inhibition of Gi/o G protein activation in the same method as described in the section of GloSensor cAMP assay.

## Acknowledgement

We thank Dr. Robert S. Molday (University of British Columbia) for kindly supplying rho 1D4-producing hybridoma. We thank Dr. David Farrens (Oregon Health & Science University) for kindly providing us with COS-1 cell line and are also grateful to Dr. Hisao Tsukamoto (Kobe University) for technical guidance on the maintenance and transfection of COS-1 cells. This work was supported by the Japanese Ministry of Education, Culture, Sports, Science and Technology Grants-in-Aid for Scientific Research JP20J01841 (YS), 23K05717 (to YI), JP24KJ1311 (to SI), 21H04969 (to HK), 22H02663 (to MK), and 23H02516 (to AT); Japan Science and Technology Agency (JST) Core Research for Evolutional Science and Technology (CREST) Grant JPMJCR1753 (to AT) and (PRESTO) grant JPMJPR19G4 (to KK).

## Author contributions

YS, MK and AT conceived and designed the study; YS, TS, YK, TO, EO and MI performed and analyzed spectroscopic experiments, HPLC and live cell assays; YI and TY were involved in high-speed time-resolved spectroscopy and YI played a leading role in conducting the SVD and global fitting analysis; SI, YT, and KK performed low-temperature UV-visible spectroscopy and analyzed results under the supervision of KK and HK; Original draft was written by YS and AT with inputs from YI, SI, and KK; All authors reviewed and gave final approval for the manuscript.

## Competing interest

The authors declare that they have competing interests.

## Notes

### Competing Interest Statement

The authors have declared no competing interest.

## References

1. A. Terakita, The opsins. Genome Biol. 6, 213 (2005).

2. A. Terakita, E. Kawano-Yamashita, M. Koyanagi, Evolution and diversity of opsins. Wiley Interdiscip. Rev. Membr. Transp. Signal. 1, 104–111 (2012).

3. M. Koyanagi, A. Terakita, Diversity of animal opsin-based pigments and their optogenetic potential. Biochim. Biophys. Acta 1837, 710–716 (2014).

4. Y. Shichida, T. Matsuyama, Evolution of opsins and phototransduction. Philos. Trans. R. Soc. Lond. B Biol. Sci. 364, 2881–2895 (2009).

5. P. D. Kiser, M. Golczak, K. Palczewski, Chemistry of the retinoid (visual) cycle. Chem. Rev. 114, 194–232 (2014).

6. A. Terakita, et al., Expression and comparative characterization of Gq-coupled invertebrate visual pigments and melanopsin. J. Neurochem. 105, 883–890 (2008).

7. T. Nagata, et al., The counterion–retinylidene Schiff base interaction of an invertebrate rhodopsin rearranges upon light activation. Commun. Biol. 2, 180 (2019).

8. R. Sato, A. Terakita, M. Koyanagi, Dragonfly red opsins share a common tuning mechanism with mammalian red opsins and further enhancement of near-infrared sensitivity. Cell. Mol. Life Sci. 83, 66 (2026).

9. T. Matsuyama, T. Yamashita, Y. Imamoto, Y. Shichida, Photochemical properties of mammalian melanopsin. Biochemistry 51, 5454–5462 (2012).

10. M. Koyanagi, K. Kubokawa, H. Tsukamoto, Y. Shichida, A. Terakita, Cephalochordate melanopsin: evolutionary linkage between invertebrate visual cells and vertebrate photosensitive retinal ganglion cells. Curr. Biol. 15, 1065–1069 (2005).

11. M. Koyanagi, et al., Bistable UV pigment in the lamprey pineal. Proc. Natl. Acad. Sci. U. S. A. 101, 6687–6691 (2004).

12. M. Koyanagi, et al., Diversification of non-visual photopigment parapinopsin in spectral sensitivity for diverse pineal functions. BMC Biol. 13, 73 (2015).

13. Y. Kakeyama, Y. Sakai, T. Sugihara, M. Koyanagi, A. Terakita, Cnidopsins characterized as bistable opsins from a reef-building coral, Acropora tenuis. Zoolog. Sci. 42, 410–416 (2025).

14. K. Sato, et al., Opn5L1 is a retinal receptor that behaves as a reverse and self-regenerating photoreceptor. Nat. Commun. 9, 1–10 (2018).

15. K. Sakai, Y. Shichida, Y. Imamoto, T. Yamashita, Creation of photocyclic vertebrate rhodopsin by single amino acid substitution. Elife 11, e75979 (2022).

16. C. Fujiyabu, et al., Amino acid residue at position 188 determines the UV-sensitive bistable property of vertebrate non-visual opsin Opn5. Communications Biology 5, 1–9 (2022).

17. Y. Sakai, et al., Coral anthozoan-specific opsins employ a novel chloride counterion for spectral tuning. Elife 14 (2025).

18. K. L. Barott, M. E. Barron, M. Tresguerres, Identification of a molecular pH sensor in coral. Proc. R. Soc. Lond. B Biol. Sci. 284, 20171769 (2017).

19. B. W. Vought, A. Dukkipatti, M. Max, B. E. Knox, R. R. Birge, Photochemistry of the primary event in short-wavelength visual opsins at low temperature. Biochemistry 38, 11287–11297 (1999).

20. A. Inoue, et al., Illuminating G-Protein-Coupling Selectivity of GPCRs. Cell 177, 1933–1947.e25 (2019).

21. N. Dascal, Signalling via the G protein-activated K+ channels. Cell. Signal. 9, 551–573 (1997).

22. K. K. Touhara, R. MacKinnon, Molecular basis of signaling specificity between GIRK channels and GPCRs. Elife 7 (2018).

23. A. Bondar, J. Lazar, Dissociated GαGTP and Gβγ protein subunits are the major activated form of heterotrimeric Gi/o proteins. J. Biol. Chem. 289, 1271–1281 (2014).

24. S. Herlitze, et al., Modulation of Ca2+ channels by G-protein beta gamma subunits. Nature 380, 258–262 (1996).

25. A. C. Dolphin, G protein modulation of voltage-gated calcium channels. Pharmacol. Rev. 55, 607–627 (2003).

26. G. W. Zamponi, K. P. M. Currie, Regulation of Ca(V)2 calcium channels by G protein coupled receptors. Biochim. Biophys. Acta 1828, 1629–1643 (2013).

27. Y. Shen, M. A. F. Rampino, R. C. Carroll, S. Nawy, G-protein-mediated inhibition of the Trp channel TRPM1 requires the Gβγ dimer. Proc. Natl. Acad. Sci. U. S. A. 109, 8752–8757 (2012).

28. J.-P. Jeon, et al., The specific activation of TRPC4 by Gi protein subtype. Biochem. Biophys. Res. Commun. 377, 538–543 (2008).

29. F. Schneider, C. Grimm, P. Hegemann, Biophysics of channelrhodopsin. Annu. Rev. Biophys. 44, 167–186 (2015).

30. M. Nack, et al., The DC gate in Channelrhodopsin-2: crucial hydrogen bonding interaction between C128 and D156. Photochem. Photobiol. Sci. 9, 194–198 (2010).

31. O. Volkov, et al., Structural insights into ion conduction by channelrhodopsin 2. Science 358 (2017).

32. A. Berndt, O. Yizhar, L. A. Gunaydin, P. Hegemann, K. Deisseroth, Bi-stable neural state switches. Nat. Neurosci. 12, 229–234 (2009).

33. O. Yizhar, et al., Neocortical excitation/inhibition balance in information processing and social dysfunction. Nature 477, 171–178 (2011).

34. C. Bamann, R. Gueta, S. Kleinlogel, G. Nagel, E. Bamberg, Structural guidance of the photocycle of channelrhodopsin-2 by an interhelical hydrogen bond. Biochemistry 49, 267–278 (2010).

35. J. Abramson, et al., Accurate structure prediction of biomolecular interactions with AlphaFold 3. Nature 630, 493–500 (2024).

36. V. A. Lórenz-Fonfría, et al., Transient protonation changes in channelrhodopsin-2 and their relevance to channel gating. Proc. Natl. Acad. Sci. U. S. A. 110, E1273–81 (2013).

37. H. Tsukamoto, Y. Kubo, A self-inactivating invertebrate opsin optically drives biased signaling toward Gβγ-dependent ion channel modulation. Proceedings of the National Academy of Sciences 120, e2301269120 (2023).

38. S. Leemann, S. Kleinlogel, Functional optimization of light-activatable Opto-GPCRs: Illuminating the importance of the proximal C-terminus in G-protein specificity. Front Cell Dev Biol 11, 1053022 (2023).

39. J. Rodgers, et al., Using a bistable animal opsin for switchable and scalable optogenetic inhibition of neurons. EMBO Rep. 22, e51866 (2021).

40. J. Wietek, et al., A bistable inhibitory optoGPCR for multiplexed optogenetic control of neural circuits. Nat. Methods 1–13 (2024).

41. M. Koyanagi, et al., High-performance optical control of GPCR signaling by bistable animal opsins MosOpn3 and LamPP in a molecular property-dependent manner. Proc. Natl. Acad. Sci. U. S. A. 119, e2204341119 (2022).

42. H. Hagio, et al., Optogenetic manipulation of Gq- and Gi/o-coupled receptor signaling in neurons and heart muscle cells. Elife 12 (2023).

43. D. T. Hartong, E. L. Berson, T. P. Dryja, Retinitis pigmentosa. Lancet 368, 1795–1809 (2006).

44. S. R. De Silva, A. T. Moore, Optogenetic approaches to therapy for inherited retinal degenerations. J. Physiol. 600, 4623–4632 (2022).

45. Q. Lu, Z.-H. Pan, Optogenetic strategies for vision restoration. Adv. Exp. Med. Biol. 1293, 545–555 (2021).

46. A. Chaffiol, J. Duebel, Mini-review: Cell type-specific optogenetic vision restoration approaches. Adv. Exp. Med. Biol. 1074, 69–73 (2018).

47. S. Nawy, The metabotropic receptor mGluR6 may signal through G(o), but not phosphodiesterase, in retinal bipolar cells. J. Neurosci. 19, 2938–2944 (1999).

48. C. Koike, et al., TRPM1 is a component of the retinal ON bipolar cell transduction channel in the mGluR6 cascade. Proc. Natl. Acad. Sci. U. S. A. 107, 332–337 (2010).

49. Y. Xu, et al., mGluR6 deletion renders the TRPM1 channel in retina inactive. J. Neurophysiol. 107, 948–957 (2012).

50. M. H. Berry, et al., Restoration of high-sensitivity and adapting vision with a cone opsin. Nat. Commun. 10, 1221 (2019).

51. J. Kralik, M. van Wyk, N. Stocker, S. Kleinlogel, Bipolar cell targeted optogenetic gene therapy restores parallel retinal signaling and high-level vision in the degenerated retina. Commun. Biol. 5, 1116 (2022).

52. M. van Wyk, S. Kleinlogel, A visual opsin from jellyfish enables precise temporal control of G protein signalling. Nat. Commun. 14, 2450 (2023).

53. J. Cehajic-Kapetanovic, et al., Restoration of vision with ectopic expression of human rod opsin. Curr. Biol. 25, 2111–2122 (2015).

54. T. Maeda, M. Golczak, A. Maeda, Retinal photodamage mediated by all-trans-retinal. Photochem. Photobiol. 88, 1309–1319 (2012).

55. A. Maeda, et al., Involvement of all-trans-retinal in acute light-induced retinopathy of mice. J. Biol. Chem. 284, 15173–15183 (2009).

56. M. Różanowska, K. Handzel, M. E. Boulton, B. Różanowski, Cytotoxicity of all-trans-retinal increases upon photodegradation. Photochem. Photobiol. 88, 1362–1372 (2012).

57. S. Levy, et al., A stony coral cell atlas illuminates the molecular and cellular basis of coral symbiosis, calcification, and immunity. Cell 184, 2973–2987 (2021).

58. S. Kayada, O. Hisatomi, F. Tokunaga, Cloning and expression of frog rhodopsin cDNA. Comp. Biochem. Physiol. B Biochem. Mol. Biol. 110, 599–604 (1995).

59. K. D. Ridge, N. G. Abdulaev, Folding and assembly of rhodopsin from expressed fragments. Methods Enzymol. 315, 59–70 (2000).

60. A. Sinha, A. M. Jones Brunette, J. F. Fay, C. T. Schafer, D. L. Farrens, Rhodopsin TM6 can interact with two separate and distinct sites on arrestin: evidence for structural plasticity and multiple docking modes in arrestin-rhodopsin binding. Biochemistry 53, 3294–3307 (2014).

61. K. Obayashi, R. Zou, T. Kawaguchi, T. Mori, H. Tsukamoto, Molecular basis underlying the specificity of an antagonist AA92593 for mammalian melanopsins. J. Biol. Chem. 301, 108461 (2025).

62. A. Terakita, R. Hara, T. Hara, Retinal-binding protein as a shuttle for retinal in the rhodopsin-retinochrome system of the squid visual cells. Vision Res. 29, 639–652 (1989).

63. Y.-C. Shen, et al., Red-tuning of the channelrhodopsin spectrum using long conjugated retinal analogues. Biochemistry 57, 5544–5556 (2018).

64. S. Inukai, K. Katayama, M. Koyanagi, A. Terakita, H. Kandori, Investigating the mechanism of photoisomerization in jellyfish rhodopsin with the counterion at an atypical position. J. Biol. Chem. 299, 104726 (2023).

65. T. Sugihara, T. Nagata, B. Mason, M. Koyanagi, A. Terakita, Absorption characteristics of vertebrate non-visual opsin, Opn3. PLoS One 11, e0161215 (2016).

66. M. Koyanagi, E. Takada, T. Nagata, H. Tsukamoto, A. Terakita, Homologs of vertebrate Opn3 potentially serve as a light sensor in nonphotoreceptive tissue. Proc. Natl. Acad. Sci. U. S. A. 110, 4998–5003 (2013).

67. M. Iwasaki, et al., Characterization of anthozoan-specific opsins from a reef-building coral, *Acropora tenuis*, as Gq-coupled opsins. Zoolog. Sci. 42, 196–205 (2025).

68. R. Matsuo, et al., Functional characterization of four opsins and two G alpha subtypes co-expressed in the molluscan rhabdomeric photoreceptor. BMC Biol. 21, 291 (2023).

