## Supplementary table and figures for "Self-regenerating opsin from reef-building coral"

### Supplementary figures

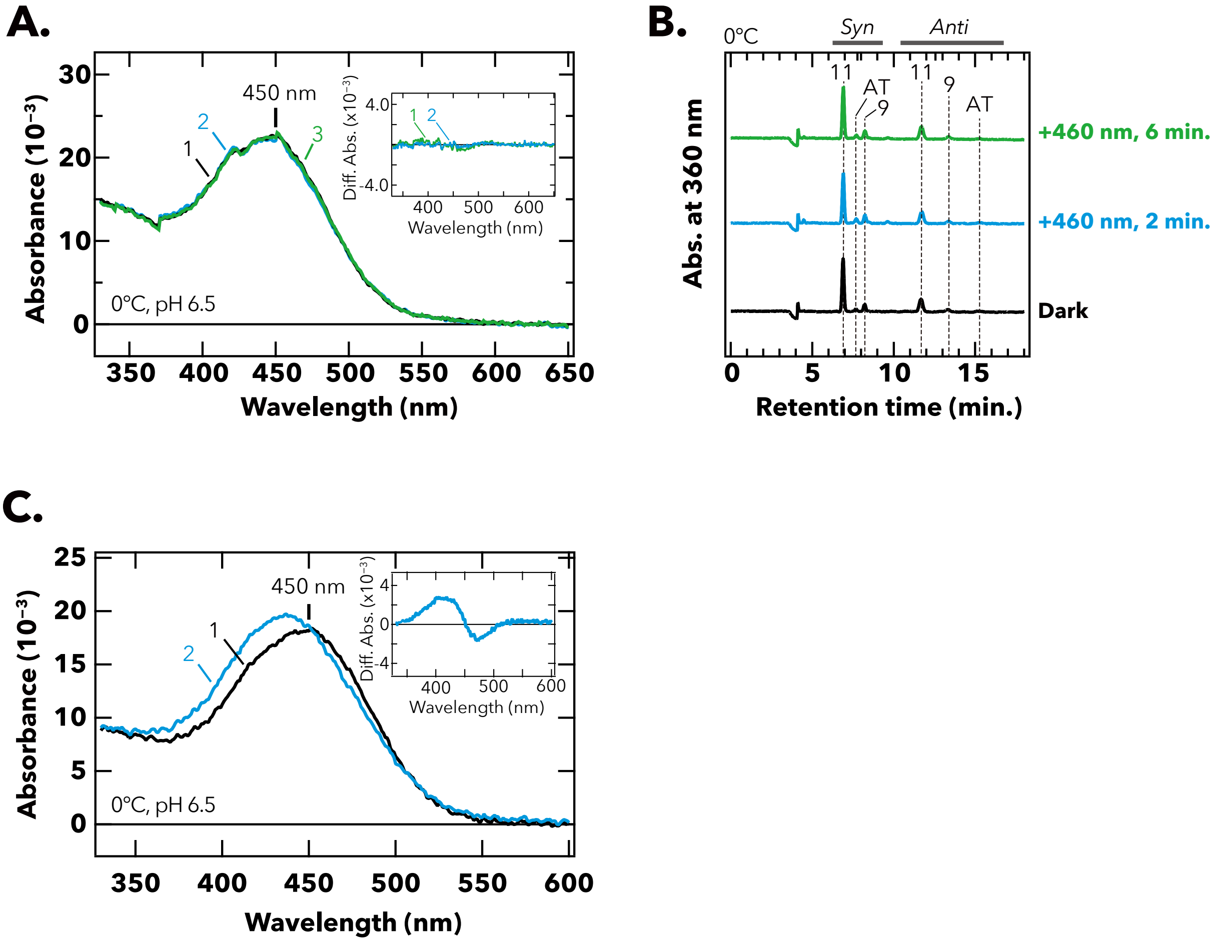


**Fig. S1. Absorption spectra and retinal configurations of wild-type and C188S-mutant AtAntho2c.** (A) Absorption spectra of wild-type AtAntho2c in the dark (curve 1) and after 2-min (curve 2) or 6-min irradiation (curve 3) with 460-nm light measured at 0°C. (*insert*) Difference spectra of before minus after 2-min- (curve 1) or 4-min-irradiation (curve 2) with blue light. (B) Isomeric configurations of retinal extracted from wild-type AtAntho2c measured by HPLC analysis. The HPLC was carried out for retinal extracted from samples before (black) and after 2-min (cyan) or 6-min irradiation (green) with 460-nm light. The samples were kept at 0°C until extracting retinal. (C) Absorption spectra of C188S-mutant AtAntho2c in the dark (curve 1) and after 1-min irradiation with 460-nm light (curve 2) measured at 0°C. (insert) Difference spectra of before minus after 1-min-irradiation with 460-nm light.

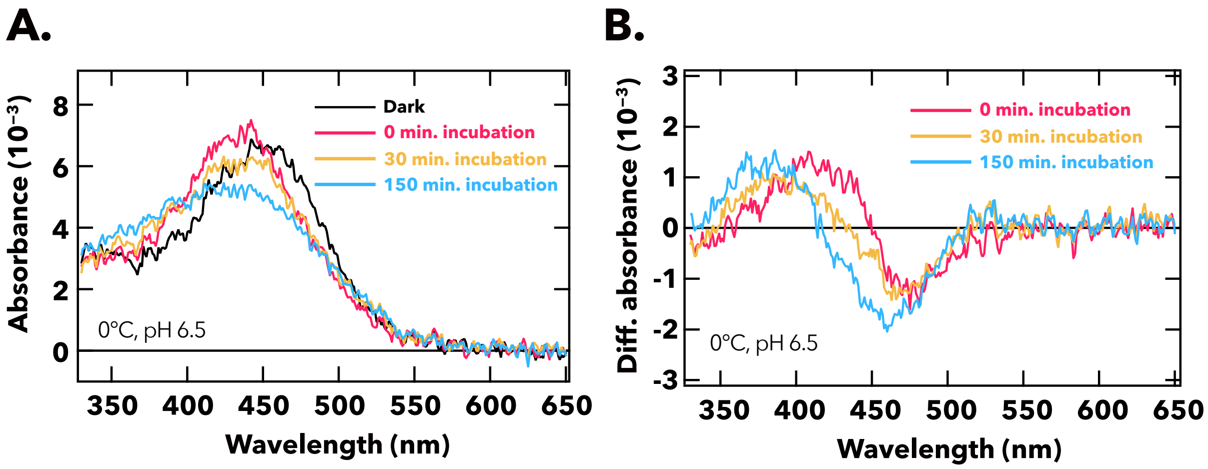


**Fig. S2. Spectral changes of C188A-mutant AtAntho2c after irradiation with yellow (> 490 nm) light.** (A) Absorption spectra of C188A-mutant AtAntho2c at 0°C and pH 6.5. The spectra were measured for the sample in the dark and from samples incubated for 0, 30, and 150 min in the dark after the light irradiation. (B) Difference spectra obtained by subtracting the spectrum in the dark (black curve in the panel A) from each spectrum measured after incubation.

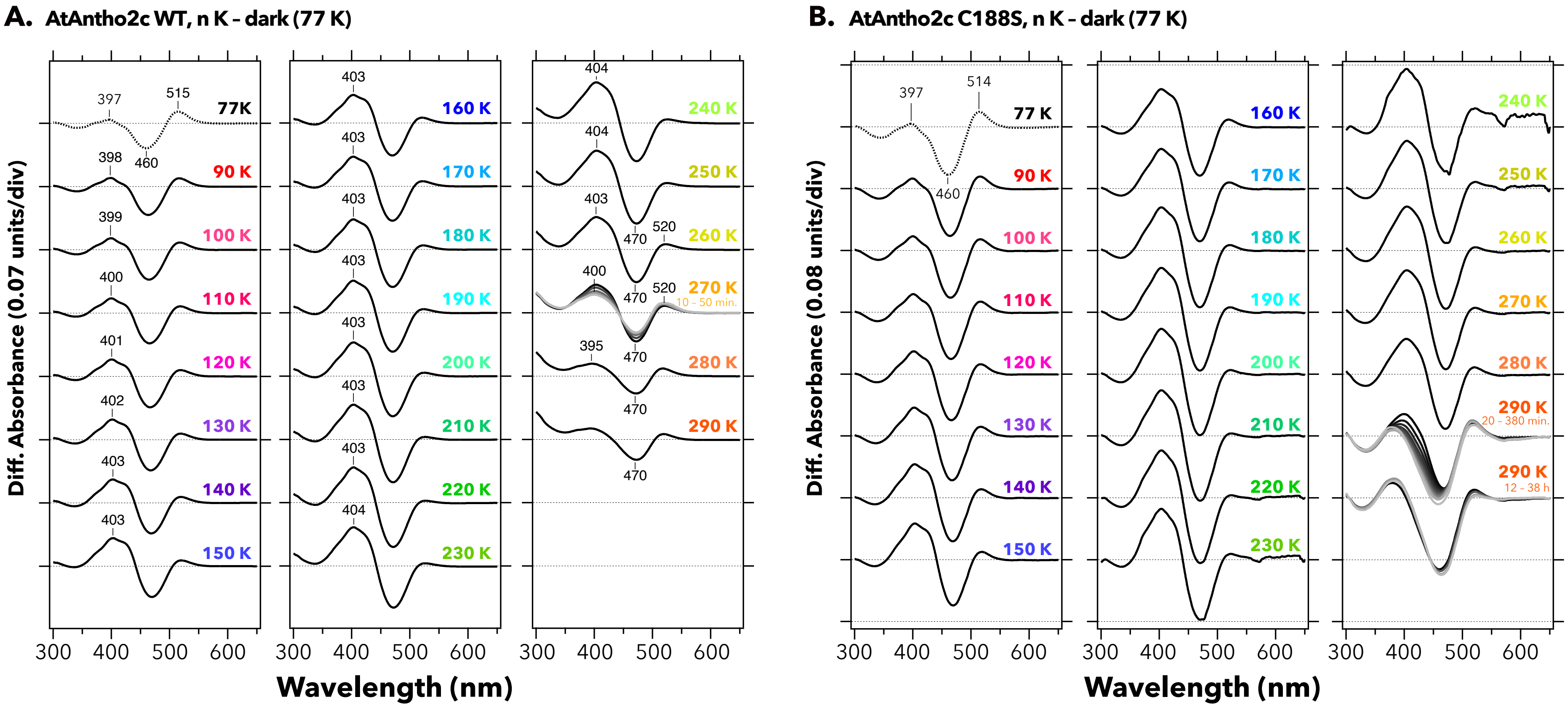


**Fig. S3. Temperature-dependent spectral changes of (A) wild-type and (B) C188S-mutant AtAntho2c after light irradiation.** UV-visible difference spectra under each different temperature condition after light irradiation against the dark spectrum as the baseline (n K – dark 77 K) are shown. Lines with different brightness at 270 K (in panel A) and 290 K (in panel B) indicate incubation times at the given temperature (darker to lighter colors correspond to longer incubation).

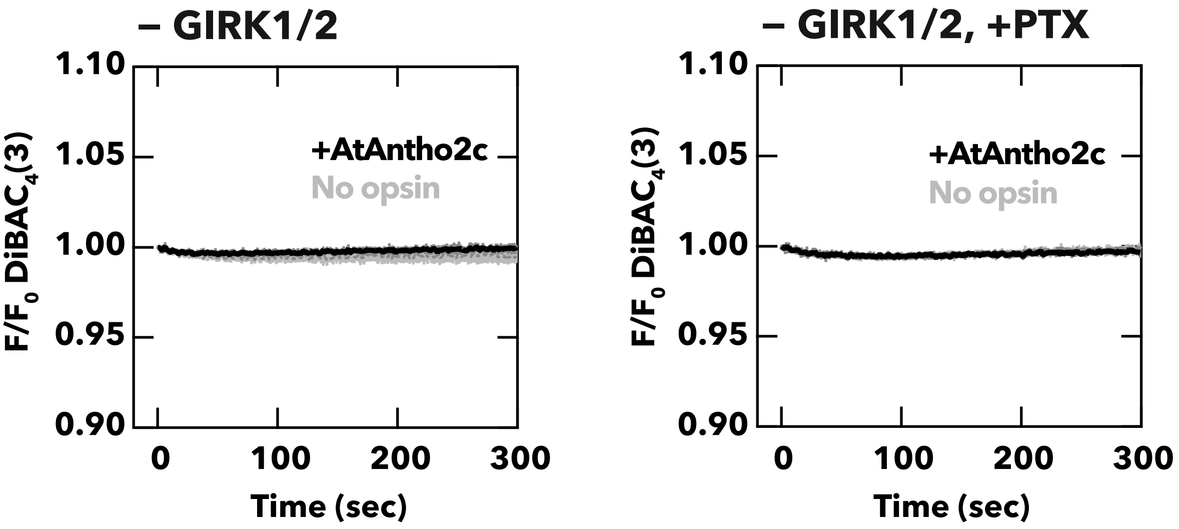


**Fig. S4. Time courses of membrane potential changes in AtAntho2c-expressing cells without GIRK1/2 co-transfection.** Normalized fluorescence levels from a membrane potential-sensitive fluorescent dye, DiBAC$_4$(3), were measured in HEK293S cells transfecting AtAntho2c (black curves) or empty vector (grey curve) without co-expression of GIRK1 and GIRK2. The cells were treated in the absence (*left* panel) or the presence (*right* panel) of pertussis toxin (PTX). Data are presented as mean (solid lines) ± standard error of the mean (shadings) for n = 3 replicates.

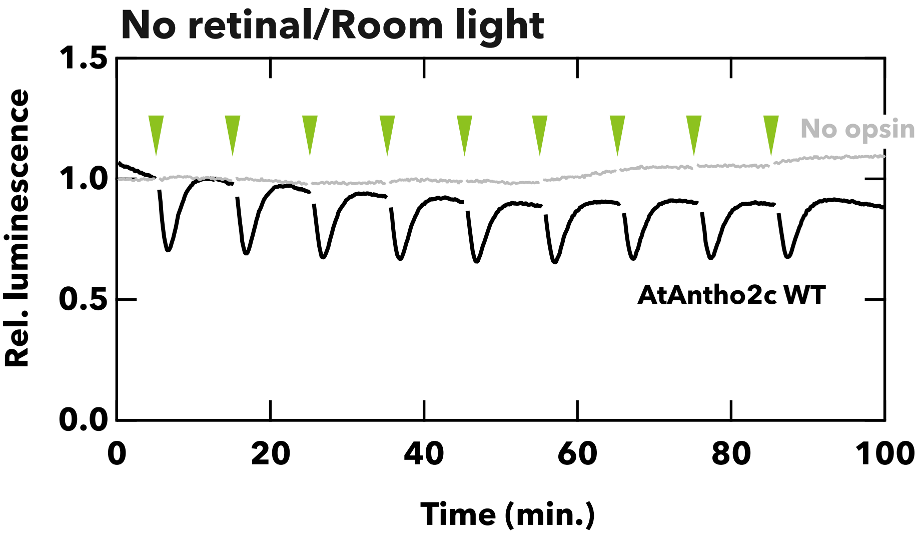


**Fig. S5. Time-course kinetics of the cAMP responses to repeated light stimulations in wild-type AtAntho2c-expressing cells treated under the room light condition.** Data from no-opsin-expressing cells was included as a negative control (grey). Transfected cells were prepared without addition of exogenous retinal and were exposed to the room light for 5 h before the recordings. It should be noted that the culture medium contains a small amount of serum-derived retinal. Green arrowheads indicate the time points at which 5-s green-light stimulation was applied. The luminescence values were normalized to the values just before the first light stimulation (time point = 0).

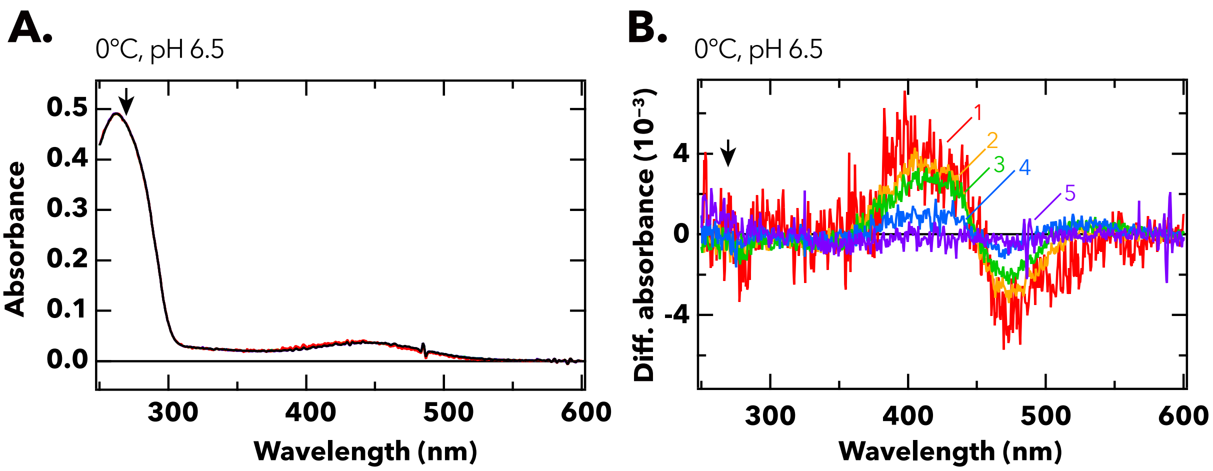


**Fig. S6. UV-visible absorption spectra and difference spectra of wild-type AtAntho2c recorded under the conditions of 0°C and pH 6.5.** (A) Absorption spectra of wild-type AtAntho2c before and after irradiation with yellow flash light. (B) Difference spectra of before minus after the light irradiation. Colors and curve numbers indicate times of the incubation in the dark after the irradiation: 1 (curve 1), 1101 (curve 2), 10101 (curve 3), 100101 (curve 4), and 631060 ms (curve 5). Black arrows highlight the absorbance at 270 nm.
